# Uterine-specific SIRT1 deficiency confers premature uterine aging and impairs invasion and spacing of blastocyst and stromal cell decidualization in mice

**DOI:** 10.1101/2021.10.25.465671

**Authors:** Magdalina J. Cummings, Hongyao Yu, Sudikshya Paudel, Guang Hu, Xiaoling Li, Myriam Hemberger, Xiaoqiu Wang

**Author notes:** To whom correspondence should be addressed.,Box 7621, 231C Polk Hall 120 W Broughton Dr. Raleigh, NC 27695-7621.

## Abstract

A distinct age-related alteration in the uterine environment has recently been identified as a prevalent cause of the reproductive decline in older female mice. However, the molecular mechanisms that underlie age-associated uterine adaptability to pregnancy are not known. Sirtuin 1 (SIRT1), a multifunctional NAD^+^-dependent deacetylase that regulates cell viability, senescence and inflammation during aging, is reduced in aged decidua. Thus, we hypothesize that SIRT1 plays a critical role in uterine adaptability to pregnancy and that uterine-specific ablation of *Sirt1* gene accelerates premature uterine aging. Female mice with uterine ablation of *Sirt1* gene using progesterone receptor Cre (*Pgr^Cre^*) exhibit subfertility and signs of premature uterine aging. These *Sirt1*-deficient mothers showed decreases in litter size from their 1^st^ pregnancy and became sterile (25.1±2.5 weeks of age) after giving birth to the 3^rd^ litter. We report that uterine-specific *Sirt1* deficiency impairs invasion and spacing of blastocysts, and stromal cell decidualization, leading to abnormal placentation. We found that these problems traced back to the very early stages of hormonal priming of the uterus. During the window of receptivity, *Sirt1* deficiency compromises uterine epithelial-stromal crosstalk, whereby estrogen (E2), progesterone (P4) and Indian hedgehog (IHH) signaling pathways are dysregulated, hampering stromal cell priming for decidualization. Uterine transcriptomic analyses also link these causes to perturbations of histone proteins and epigenetic modifiers, as well as adrenomedullin signaling, hyaluronic acid metabolism, and cell senescence. Strikingly, our results also identified genes with significant overlaps with the transcriptome of uteri from aged mice and transcriptomes related to master regulators of decidualization (e.g., *Foxo1, Wnt4, Sox17, Bmp2, Egfr* and *Nr2f2*). Our results also implicate accelerated deposition of aging-related fibrillar type I and III collagens in *Sirt1*-deficient uteri. Collectively, SIRT1 is an indispensable age-related regulator of invasion and spacing of blastocysts, as well as decidualization of stromal cells.

**Author Summary:** Advanced maternal age (i.e., ≥35 years old) is considered a significant risk factor for birth defects, including fetal growth restriction, stillbirth, preterm birth, and preeclampsia. Lifestyle factors such as tobacco smoke, alcohol usage and other environmental toxins are facilitators of premature reproductive aging. Thus, understanding the precise mechanisms by which female reproductive organs age is a prerequisite for ultimately developing counteracting therapies. Much attention has been focused on ovarian function and oocyte quality, but we provide evidence that severe placentation defects are a major cause of age-related reproductive decline, which results from an impaired decidual response by uterine stromal cells. This problem is due to a blunted progesterone (P4) responsiveness of the aging uterus, via its cognate receptor, PGR. In the present study, the uterine SIRT1 gene was deleted to advance understanding of the genetics of premature uterine aging in a mouse model of blunted PGR actions that is similar to physiological aging. Our results have informed as to molecular changes in response to blunting PGR actions in aging uteri that are unrelated to oocyte quality, and provide new insights into strategies for developing counteracting measures for pregnancy in females at an advanced reproductive age.

## Introduction

Advanced maternal age (i.e., ≥35 years old) is a major risk factor for birth defects [1–6], which occur in 3-5% of pregnancies [7]. These adverse pregnancy outcomes include, but are not limited to miscarriage, late fetal and perinatal death, stillbirth, preterm birth and extreme preterm birth, preeclampsia, and intrauterine growth restriction, as well as maternal medical complications such as gestational diabetes and maternal mortality [5, 8–14]. In women over 40 years of age, the incidence of spontaneous abortion can increase to >30% [15]. Thus, understanding the precise mechanisms by which female reproductive organs age is a prerequisite for ultimately developing counteracting measures. Much attention has been focused on ovarian function and oocyte quality [6, 16], but recent evidence has shown that uterine decidualization and placentation defects are a major cause of age-related declines in fertility [17]. However, the underlying mechanisms by which reproductive aging affects uterine adaptability to pregnancy are largely unknown.

Sirtuin 1 (SIRT1) is a nicotinamide adenosine dinucleotide (NAD^+^)-dependent histone deacetylase (HDAC) that acts on epigenetic and non-epigenetic targets, and also regulate transcription machinery binding to specific chromosomal regions [18–21]. The molecular actions of SIRT1 regulate cell cycle, DNA repair, adipogenesis, glucose output, insulin sensitivity, fatty acid oxidation, neurogenesis, apoptosis and oxidative stress, thereby affecting cell viability, senescence, inflammation during aging [18, 22–30]. In particular, SIRT1 and its ortholog have a key role in delaying cell senescence and extending the lifespan of yeast, flies, and mice [31–33]. Fertility of female *Sirt1*-null mice is significantly compromised. *Sirt1*-null female derived from deletions of exons 5 and 6 are essentially sterile with only one female producing 3 litters within 7 months of breeding trial [34]. *Sirt1*-null female mice derived from deletion of exon 4 are subfertile with significant decreases in litter size and numbers of litters in a fixed breeding period [35]. *In vitro* studies suggest that SIRT1 interacts with cognate nuclear receptor ESR and PGR to affect E2- or P4-dependent transcriptional activity [36–38]. However, uterine-specific roles of SIRT1 during pregnancy and its roles in uterine aging are not known. Here, we show that SIRT1 decreases in the uterine decidua of aged female mice and is directly regulated by SOX17 and FOXO1, two critical co-regulators of PGR action required for implantation and decidualization [39, 40]. Thus, we hypothesize that SIRT1 plays a critical role in governing age-related uterine receptivity and uterine-specific ablation of SIRT1 accelerates premature uterine aging. Our results revealed that *Sirt1* mRNA is expressed in uterine luminal (LE) and glandular (GE) epithelia, as well as stromal cells (S) of murine endometria during early pregnancy. Female mice with uterine ablation of the *Sirt1* gene (*Sirt1^d/d^*; *Pgr^Cre/+^Sirt1^f/f^*) are subfertile and exhibit signs of premature uterine aging. These *Sirt1^d/d^* mothers experienced decreases in litter size from their 1^st^ pregnancy and became sterile (25.1±2.5 weeks of age) after giving birth to their 3^rd^ litter. We report that uterine-specific *Sirt1* deficiency impairs invasion and spacing of blastocysts, and stromal cell decidualization, leading to abnormal placentation. Moreover, we traced these *Sirt1*-associated problems to defects in epithelial-stromal crosstalk during the window of uterine receptivity to implantation, in which E2, P4, and IHH cell signaling pathway are dysregulated, hampering stromal cell priming for decidualization. Further transcriptomic analyses not only substantiated dysregulations of E2, P4 and IHH pathways in *Sirt1*-deficient uteri, but also linked the causes to perturbations of histone proteins and epigenetic modifiers, as well as pathways including adrenomedullin and hyaluronic acid signaling, and cell senescence. We then demonstrated that the SIRT1-deficient transcriptome is highly similar to that of mice with an aged uterine transcriptome. Accelerated deposition of aging-related fibrillar type I and III collagens in *Sirt1*- deficient uteri further suggests that SIRT1 is the indispensable age-associated regulator for implantation, decidualization and placentation.

## Results

### Advanced maternal age impairs uterine decidualization

The ability of the endometrial stromal cells to undergo artificially induced decidualization was determined. The uteri of young female mice (6-8 weeks old) displayed a robust decidual response, as evidenced by increased size and wet weight of the decidua formed in the stimulated right uterine horn at decidual day 2 (DD2) and 5 (DD5) (**Fig 1A and B**). However, the uteri of aged females (46-54 weeks old) failed to form a robust decidua at DD2 and DD5 (**Fig 1A**), and the ratio of decidual to control horn weight was less than that of young female mice at DD2 (*P*<0.01) and DD5 (*P*<0.001) (**Fig 1B**). H&E staining of the mid-section of uterine decidua was conducted to determine the impact of advanced maternal age on the progression of uterine decidualization. After the artificial deciduogenic stimulus at DD0, the uteri of young females exhibited increases in endometrial vascular permeability and development of stromal edema, with enlarged stromal cells with rounded nuclei beginning in the anti-mesometrial pole (AM) of the endometrium at DD2 and spreading to the mesometrial (M) stromal cells at DD5 (**Fig 1C**). On the other hand, these macroscopically identifiable events were not evident in uteri of aged females at DD2 and DD5 (**Fig 1C**). Moreover, the endometrial LE remained intact and the center of the decidual tissue in aged mice was fluid-filled at DD5.

**Fig 1.**
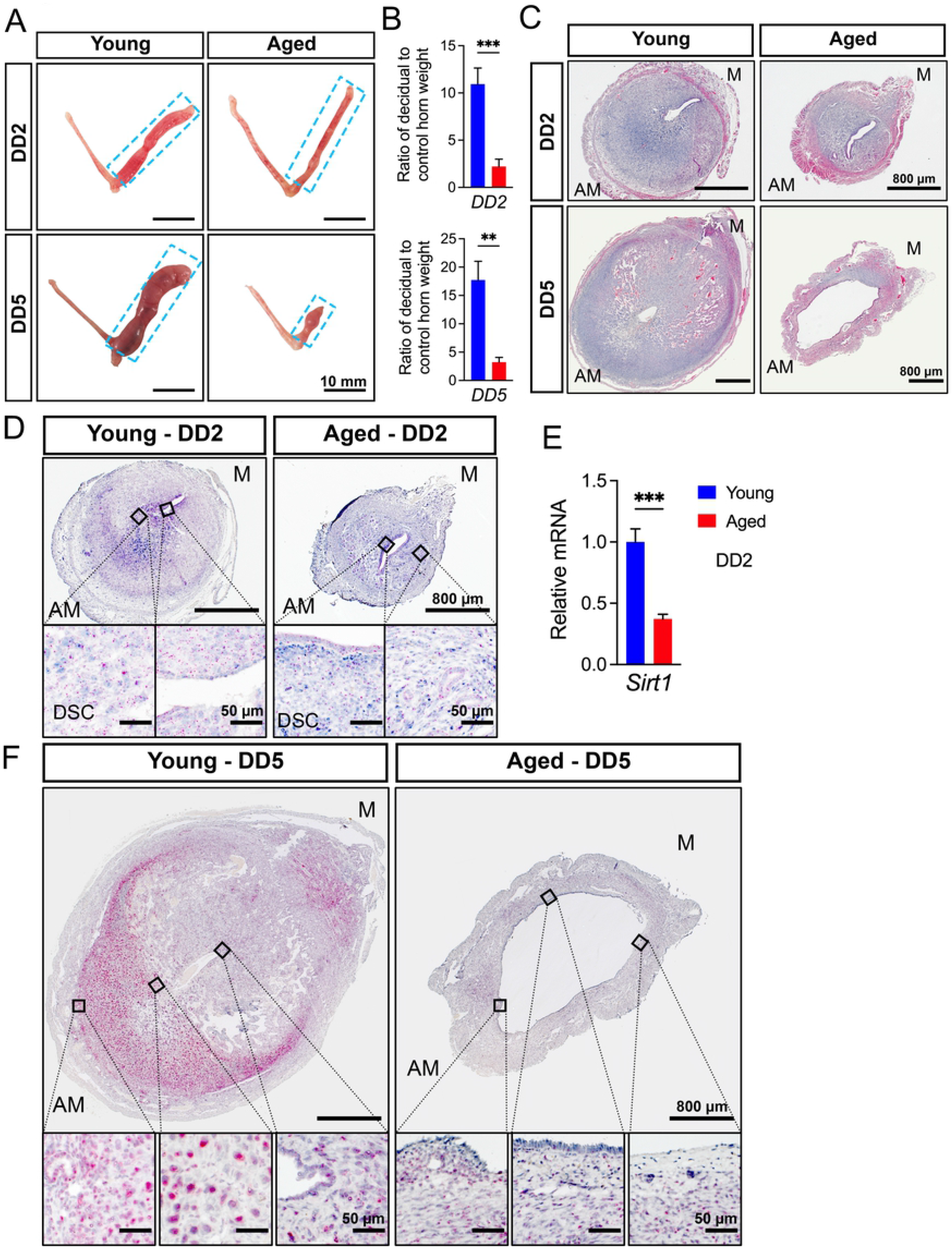
Advanced maternal age impacts artificial decidua formation and *Sirt1* mRNA expression. **(A)** Artificial decidua formation in uteri of young and aged C57BL/6 females at DD2 and DD5. Blue dashed squares denote decidual tissue in the uterine horn. **(B)** Ratios of decidual uterine horn weight to control uterine horn weight in young and aged females at DD2 and DD5. **(C)** Histological analyses following H&E staining of uterine decidua in young and aged females at DD2 and DD5. **(D-F)** RNAscope *in situ* hybridization (D,F) and qRT-PCR analyses (E) of *Sirt1* mRNA in decidua in young and aged females at DD2 and/or DD5. M, mesometrial side; AM, anti-mesometrial side. n=6 for DD2; and n=4 for DD5. DD2, decidual day 2; DD5, decidual day 5. **, *P*<0.01; ***, *P*<0.001 (two-tailed *t* test). Data are presented as means ± SEM

### SIRT1 expression decreases in decidua of aged mice

Localization and expression of *Sirt1* mRNA in the endometrial compartments of artificially decidualized uteri were determined using *in situ* hybridization and qRT-PCR in young and aged female mice (**Fig 1D-F**). At DD2, expression of *Sirt1* mRNA was detectable in uterine LE, GE and S, but mainly in differentiated stromal cells (DSCs) of young and aged uteri (**Fig 1D**). Compared with robust DSCs in young uteri, *Sirt1* mRNA expression decreased in the less differentiated stromal cells of aged uteri (**Fig 1D**). The qRT-PCR analyses further demonstrated decreases (*P*<0.001) in the abundance of *Sirt1* mRNA in aged decidua at DD2 (**Fig 1E**). At DD5, *Sirt1* mRNA was expressed strongly in uterine LE, GE and DSCs, particularly anti-mesometrial DSCs in uteri of young mice; whereas its abundance was much less in decidual of aged mice (**Fig 1F**).

### Regulation of endometrial SIRT1 during the peri-implantation period of pregnancy

The temporal and cell-specific expression of *Sirt1* mRNA in the uterus on GD 0.5 to 5.5 was determined in young wildtype female mice using RNA *in situ* hybridization (**Fig 2A**). GD 0.5 corresponds to the morning of the presence of the postcoital plug. Expression of *Sirt1* mRNA was detected in both uterine LE and GE between GD 0.5 and 5.5; while stromal expression of *Sirt1* mRNA increased between GD 1.5 and 3.5. By GD 4.5, *Sirt1* mRNA was expressed in uterine LE and DSCs surrounding the implantation sites (IS) of blastocysts (E); and increased significantly in the primary (PDZ) and secondary decidual zone (SDZ) at GD 5.5. qRT-PCR analyses confirm that expression of *Sirt1* mRNA increased (*P*<0.05) more in uteri of the mice between GD 1.5 and 2.5 as compared to GD 0.5, 3.5 and PDD 4.5 (**Fig 2B**). After mining the independent mouse ChIP-seq datasets (GSE34927, GSE36455, GSE118327 and GSE72892), we identified PGR, ESR, SOX17 and FOXO1 as transcription factors sharing a common binding peak at the promoter region of the *Sirt1* gene in uteri of P4-treated, E2-treated, and uteri of pregnant mice on GD 3.5 or GD 4.5 (**Fig 2C**). To determine whether SOX17 or FOXO1 directly regulate SIRT1 in uteri of female mice, mice with uterine-specific ablation of *Sox17* (*Sox17^d/d^*) or *Foxo1* (*Foxo1^d/d^*) were evaluated. RNA *in situ* hybridization analyses showed that expression of *Sirt1* mRNA decreased 1) in uterine LE, GE and S of *Sox17^d/d^* uteri at GD 3.5 (**Fig 2D**); and 2) in LE and DSCs of *Foxo1^f/f^* uteri within the implantation chamber on GD 4.5 (**Fig 2E**).

**Fig 2.**
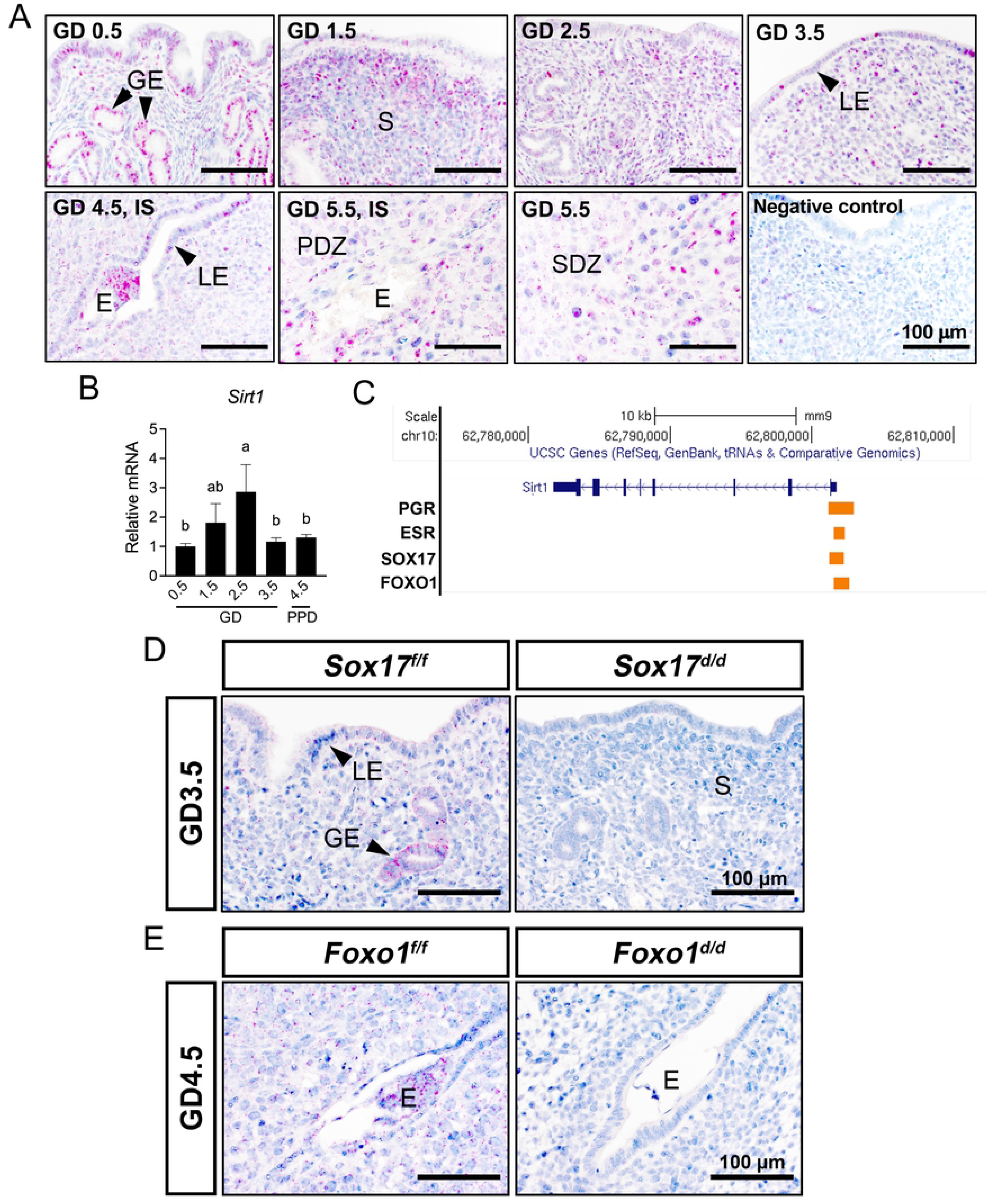
Regulation of expression of *Sirt1* mRNA in uteri of young female mice during the peri-implantation period of pregnancy. **(A)** RNAscope *in situ* hybridization for *Sirt1* mRNA in uteri of young female mice between GD 0.5 and 5.5 (n=3). **(B)** Quantification of *Sirt1* mRNA in uteri from young female mice between GD 0.5 and 3.5, as well as PPD 4.5 (n=6). **(C)** Genome Browser tracks of PGR, ESR, SOX17 and FOXO1 binding at promoter region for *Sirt1* gene in P4-treated, E2-treated, GD 3.5, or GD 4.5 uteri. UCSC Genome Browser views showing the mapped read coverage of PGR, ESR, SOX17, and FOXO1 ChIP-seq data (GSE34927. GSE36455, GSE118327, and GSE72892). **(D)** RNAscope *in situ* hybridization for *Sirt1* mRNA in uteri from *Sox17^f/f^* and *Sox17^d/d^* mice (n=3) on GD 3.5. **(E)** RNAscope *in situ* hybridization for *Sirt1* mRNA in uteri from *Foxo1^f/f^* and *Foxo1^d/d^* mice (n=3) on GD 4.5. GD, gestational day; PPD, pseudopregnant day; IS, implantation site; IIS, inter-implantation site; LE, luminal epithelium; GE, glandular epithelium; S, stroma; E, embryo; PDZ, primary decidual zone; SDZ, secondary decidual zone. Different superscript letters denote significant differences (*P*<0.05, ANOVA with Fisher’s LSD post-hoc test). Data are presented as means ± SEM

### Female mice lacking uterine SIRT1 are subfertile

The physiological role(s) of SIRT1 in uteri of adult mice was investigated by breeding mice with a conditional allele of *Sirt1* (*Sirt1^f/f^* mice) [41, 42] with *Pgr^Cre^* mice [43] to ablate *Sirt1* in PGR-positive cells within the uterus (*Sirt1^d/d^*). Analyses by qRT-PCR confirmed that expression of *Sirt1* mRNA (P<0.01) in whole uteri of *Sirt1^d/d^* females was ablated as compared with results from evaluation of uteri from *Sirt1^f/f^* mice (**Fig 3A**). Evidence of ablation of SIRT1 protein was also confirmed by Western blot analyses (**Fig 3B**).

**Fig 3.**
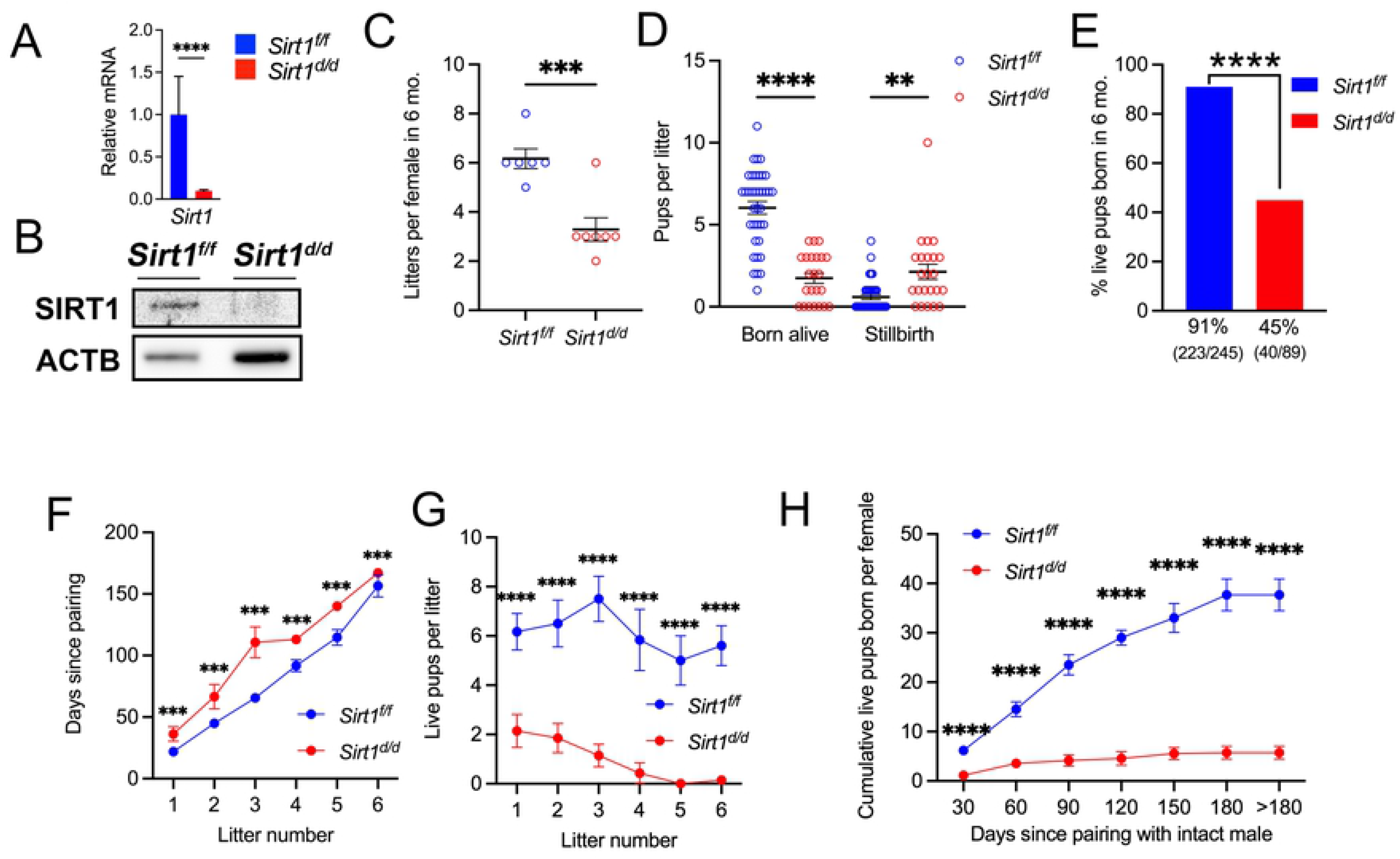
Uterine-specific deletion of *Sirt1* impacts the fertility of female mice in a 6-month breeding trial. (A-B) Generation of *Sirt1* conditional knockout (*Sirt1^d/d^*; *Pgr^Cre/+^Sirt1^f/f^*) female mice using the *Pgr^Cre^* mouse model was validated by using qRT-PCR (A) and Western blot (B) analyses. Gene expression was normalized to 18s rRNA in qRT-PCR analyses; and β-actin (ACTB) was used as protein loading control in Western blot analyses. \*\*\*\**P*<0.0001 (two-tailed *t* test). **(C)** Litters produced from *Sirt1^f/f^* (control) and *Sirt1^d/d^* female mice. Each dot represents the value for a different individual. \*\*\**P* < 0.001(two-tailed *t* test). **(D)** Stillbirth and live pups born per litter in *Sirt1^f/f^* and *Sirt1^d/d^* female mice. Each dot represents the value for a different litter. \*\**P* < 0.01; \*\*\*\**P* < 0.0001 (two-tailed *t* test). **(E)** Percent live pups born in 6 months from *Sirt1^f/f^* and *Sirt1^d/d^* female mice. \*\*\*\**P*<0.0001 (χ^2^ test). **(F)** Time taken to produce a specific number of litters. **(G)** Live pups born in a specific number of litters. **(H)** Cumulative live pups born since pairing. The mean number of pups produced up to each time point is denoted along with the respective error bar, with >180 d representing the day for calculating the total number of live pups produced. ***, *P*<0.001; ****, *P*<0.0001 (Two-way ANOVA with Tukey’s multiple comparisons test) at specific time points for contrast between *Sirt1^f/f^* (in blue, n=6) and *Sirt1^d/d^* (in red, n=7) female mice. Data are presented as means ± SEM. Also see **S1 Fig**.

*Sirt1^d/d^* female mice were subfertile. During a 6 months of breeding trial, *Sirt1^f/f^* mice produced 6.2±0.4 litters per female (n=6 females) with an average of 6.6±0.4 pups per litter (n=37 litters); whereas *Sirt1^d/d^* mice produced only 3.3±0.5 litters per female (*P*<0.001; n=7 females) with an average of 3.9±0.5 pups per litter (*P*<0.0001; n=23 litters) (**Fig 3C and S1 Fig A**). Among pups born per litter, control females gave birth to 6.0±0.4 live pups with 0.6±0.2 stillbirth; whereas *Sirt1^d/d^* females gave birth to 1.7±0.3 live pups (*P*<0.0001) with 2.1±0.5 stillbirth (*P*<0.01) (**Fig 3D**). Thus, the percentage of live pups born from *Sirt1^d/d^* mothers (223/245; 45%) in 6 months was less (*P*<0.0001) than for *Sirt1^f/f^* control mice (40/89; 91%) (**Fig 3E and S1 Fig B**). To further assess fecundity over time in females paired with intact males, we examined the frequency of production of litters of mice (i.e., the rate of litter production) (**Fig 3F**), pups per litter at specific litters (**Fig 3G and S1 Fig C**), and a total number of pups produced per female up to specific time points (i.e., cumulative pups born per female) (**Fig 3H and S1 Fig D**). A uterine SIRT1 deficiency revealed that *Sirt1^d/d^* females took a much longer time (*P*<0.001) to produce each litter as compared with *Sirt1^f/f^* mothers (**Fig 3F**) and had smaller litters (*P*<0.0001; **S1 Fig C**), fewer live pups (*P*<0.0001; **Fig 3G**), and total pups per litter from their 1^st^ pregnancy onward. Further, 6 out of 7 *Sirt1^d/d^* mothers were sterile by 25.1±2.5 weeks of age after giving birth to their 3^rd^ litter (**Fig 3F and G**). All of those indices indicate that SIRT1 is necessary to prevent premature reproductive aging. Similarly, cumulative numbers of pups born per female were greater (*P*<0.05) for *Sirt1^f/f^* than *Sirt1^d/d^* mice at 30 days after pairing, and remained different (*P*<0.0001) thereafter (**Fig 3H and S1 Fig D**). Moreover, a plateau was observed in both cumulative total (12.7±1.7) and live (5.7±1.3) pups produced per *Sirt1^d/d^* female at 150 days after pairing. On the other hand, *Sirt1^f/f^* mice continued to produce regular litters throughout the 6 months of the breeding trial, i.e., 40.8±4.0 pups as a cumulative total and 37.7±3.2 live pups as a cumulative total per female.

### SIRT1 impacts invasion and spacing of blastocysts, and decidualization of uterine stromal cells

In order to determine the cause of the subfertility in *Sirt1^d/d^* females, we determined effects on spacing, invasion and implantation of blastocysts, as well as decidualization of uterine stromal cells. Mice were killed at GD 5.5, 6.5 and 9.5 and the uteri were examined for the presence of implantation sites. As shown in **Fig 4A and B**, the number of implantation sites for blastocysts was not different between *Sirt1^f/f^* and *Sirt1^d/d^* females. However, mice with the SIRT1 deficiency had lower (*P*<0.0001) weights of the decidua at GD 6.5 and spacing of implantation sites was more uneven (*P*<0.05) on GD 9.5 (**Fig 4C-E**). H&E staining of the mid-section of implantation sites was conducted to determine the impact of SIRT1 deficiency on blastocyst invasion and progression of uterine decidualization. Compared with *Sirt1^f/f^* uteri in which blastocysts invaded towards anti-mesometrial decidual cells, blastocyst remained in the center of *Sirt1^d/d^* uteri at GD 5.5 and 6.5 (**Fig 4F, 1^st^ and 2^nd^ columns**), as demonstrated by distance ratio of the implantation sites to each side (AM or M) at the endometrial-myometrial interface (**Fig 4G**). By GD 9.5, the definitive placentas of *Sirt1^d/d^* females were defective and the anti-mesometrial decidual cells did not undergo proper apoptosis to form the decidua capsularis, insofar as the trophoblast portion was severely under-developed and the main direction of placentation was off-center (**Fig 4F, 3^rd^ column**). In addition, the ability of the endometrial stromal cells to undergo an artificially induced decidualization was assayed. The uteri of *Sirt1^f/f^* mice displayed a robust decidual response, as evidenced by increased size and wet weight of the decidua in the stimulated right uterine horn 5 (DD5) days after stimulus delivery (**Fig 4H**). However, the decidual/control weight ratio at DD5 was less (*P*<0.05) in the uteri of *Sirt1^d/d^* mice (**Fig 4H**).

**Fig 4.**
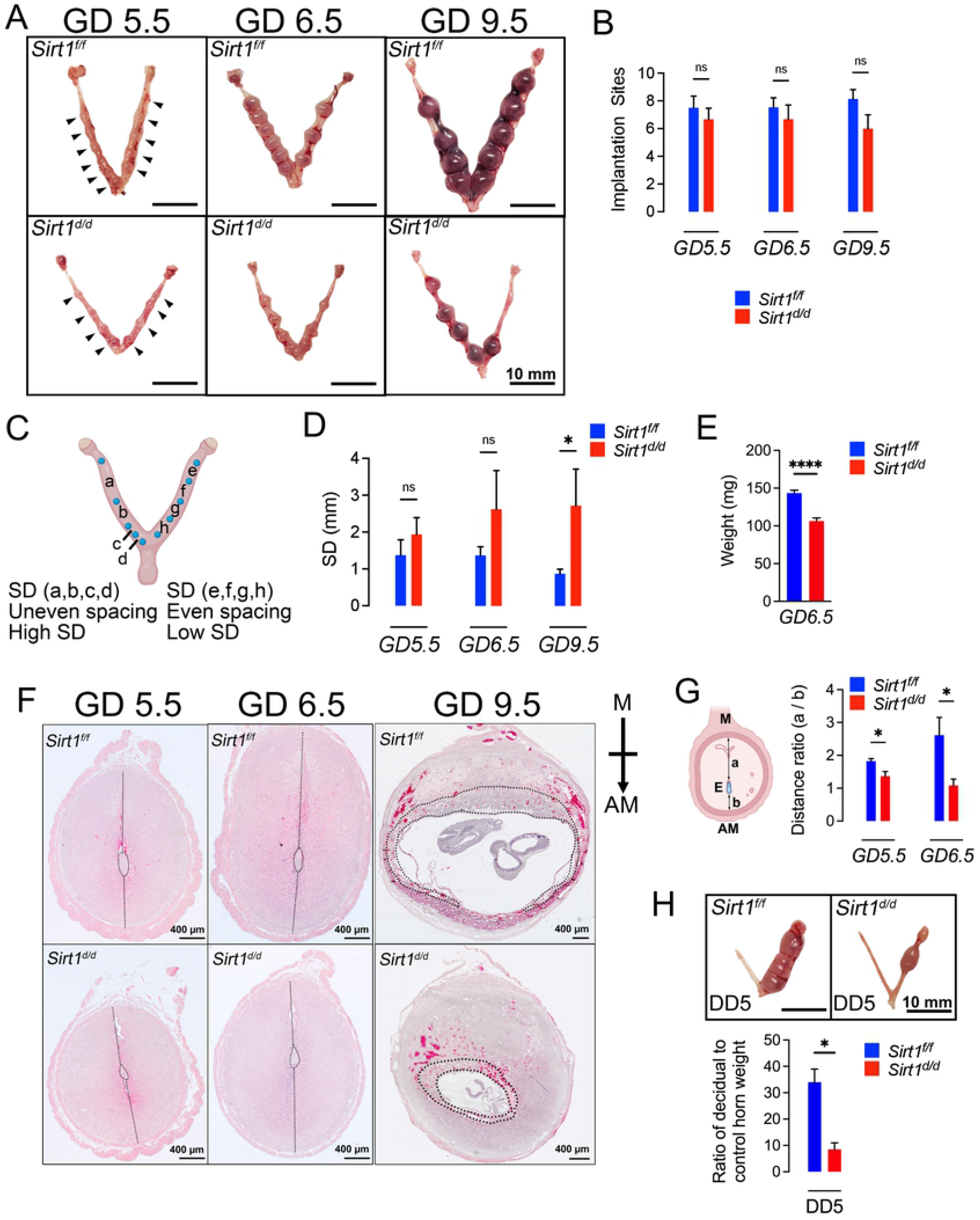
Uterine-specific deletion of *Sirt1* impacts spacing and invasion of blastocysts and stromal cell decidualization. **(A,B)** Images (A) and quantification (B) of implantation sites in *Sirt1^f/f^* and *Sirt1^d/d^* female mice (n=6). NS, not significant (two-tailed *t* test). **(C)** Illustration of quantitation method for spacing of blastocysts within uterine horns. SD, standard deviation. **(D)** Quantification of spacing of blastocysts in uteri from *Sirt1^f/f^* and *Sirt1^d/d^* females using the method depicted in (C). n=6 females per groups. \**P*<0.05 (two-tailed *t* test). **(E)** Decidual weight in *Sirt1^f/f^* and *Sirt1^d/d^* female mice (n=6). \*\*\*\**P* < 0.0001 (two-tailed *t* test). **(F)** Histological analyses following H&E staining of uteri from *Sirt1^f/f^* and *Sirt1^d/d^* female mice (n=6). Dotted lines in GD 5.5 and 6.5 indicate the distance from implantation site to AM or M; and dotted lines in GD 9.5 indicate boundary between the fetal trophoblast compartment and the maternal decidua. M, mesometrial pole; AM, anti-mesometrial pole. **(G)** Ratio of the distance between E-to-M to distance between E-to-AM in uteri from *Sirt1^f/f^* and *Sirt1^d/d^* female mice at GD 5.5 and 6.5 (n=6). E, embryo. \**P*<0.05 (two-tailed *t* test). **(H)** Artificial decidua formation and ratio of decidual to control horn weight in *Sirt1^f/f^* and *Sirt1^d/d^* female mice (n=6). \**P*<0.05 (two-tailed *t* test). Data are presented as means ± SEM

### SIRT1 dysregulates localization of PTGS2, FOXO1 and PGR in post-implantation uteri

To determine the possible causes of these decidualization defects, cell-specific expression patterns for PTGS2, FOXO1 and PGR were assessed on GD 5.5, using immunohistochemical analyses, as they are required for decidualization [39, 44, 45] (**Fig 5**). PTGS2 was highly expressed in *Sirt1^f/f^* uteri and localized to the mesometrial DSCs at sites of invading blastocysts; whereas, in *Sirt1^d/d^* uteri, PTGS2 was expressed in the DSCs closely surrounding the entire blastocyst, suggesting a delay in implantation and dysregulation of decidualization at GD 5.5 (**Fig 5A**). Differences in FOXO1 localization confirmed altered implantation as FOXO1 remained localized to nuclei of LE at the fetal-maternal interface and expression was less in mesometrial LE and S of GD 5.5 *Sirt1^d/d^* uteri (**Fig 5B**). Staining of PGR further showed that SIRT1 deficiency had strong effects on inhibition of PGR expression and stromal cell decidualization, demonstrated by downregulation of PGR in DSCs close to the blastocyst and reduced area of DSCs in the endometrium of GD 5.5 *Sirt1^d/d^* uteri.

**Fig 5.**
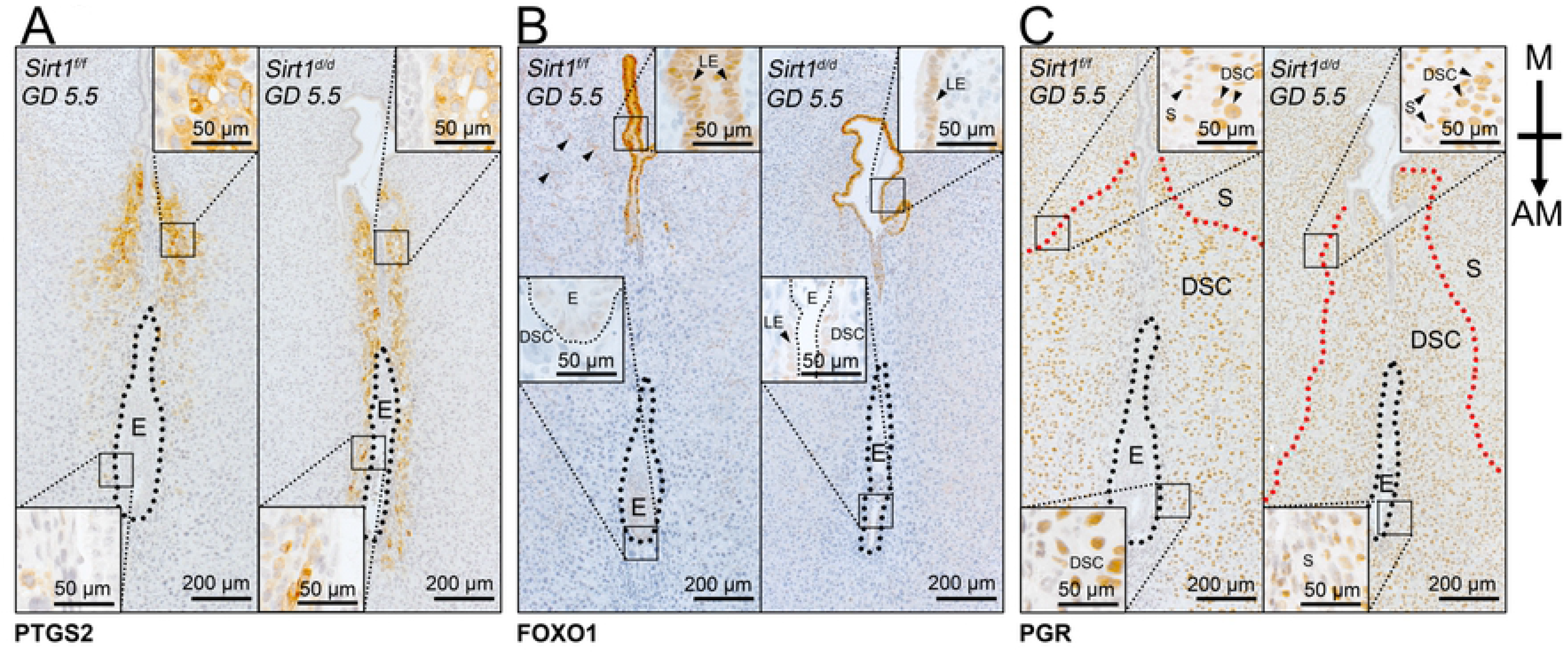
Immunohistochemical staining of PTGS2 (A), FOXO1 (B) and PGR (C) proteins in uteri from *Sirt1^f/f^* and *Sirt1^d/d^* female mice on GD 5.5. Black dotted circles indicate sites of blastocyst implantation. Red dotted lines indicate the boundary between the DSC and undifferentiated S. Inserts indicate high magnification of box area. E, blastocyst; LE, luminal epithelium; S, stroma; DSC, decidualized stromal cells; M, mesometrial pole; AM, anti- mesometrial pole.

### SIRT1 regulates uterine E2 and P4 signaling

Our next approaches were aimed at identifying differences in uterine responses during the window of receptivity to implantation when uterine epithelia stop proliferating, becomes permissive for blastocyst attachment and invasion, and primes stromal cells for decidualization. For this purpose, we mated *Sirt1^f/f^* and *Sirt1^d/d^* females with intact wildtype males and dissected uterine horns at GD 3.5 for immunohistochemical and qRT-PCR analyses. Given that uterine function is orchestrated by ovarian steroid hormones [46–49], the effect of *Sirt1* ablation on E2 and P4 signaling was investigated. Immunohistochemical analyses of the uteri from 6-8 week old *Sirt1^d/d^* mice on GD 3.5, showed increases in estrogen receptor (ESR1) in epithelial cells (**Fig 6A, top panel**), but no difference in expression of *Esr1* mRNA was detected between *Sirt1^f/f^* and *Sirt1^d/d^* uteri (**Fig 6B, top panel**). Concomitantly, as downstream targets of ESR1, *Ltf* mRNA expression was not different, but *Lif* and *Lifr* expression was lower (*P*<0.05) in *Sirt1^d/d^* uteri as compared to *Sirt1^f/f^* control at GD 3.5 (**Fig 6B, top panel**). Progesterone receptors (PGR) were reduced in both epithelial cells and stromal cells of *Sirt1^d/d^* uteri (**Fig 6A, bottom panel**). The expression of *Pgr* mRNA, as well as mRNAs for its downstream targets, *Areg* and *Ihh* were also downregulated (*P*<0.01) in the *Sirt1^d/d^* uteri (**Fig 6B, 2^nd^ panel**). As co-regulators for PGR action, expression of *Foxo1* and *Arid1a mRNAs* was downregulated in the *Sirt1^d/d^* uteri; and there was no difference in expression of *Sox17* between *Sirt1^f/f^* and *Sirt1^d/d^* uteri (**Fig 6B, bottom panel**). In addition, expression of *Foxa2* protein and mRNA was not different between *Sirt1^f/f^* and *Sirt1^d/d^* uteri (**S2 Fig**).

**Fig 6.**
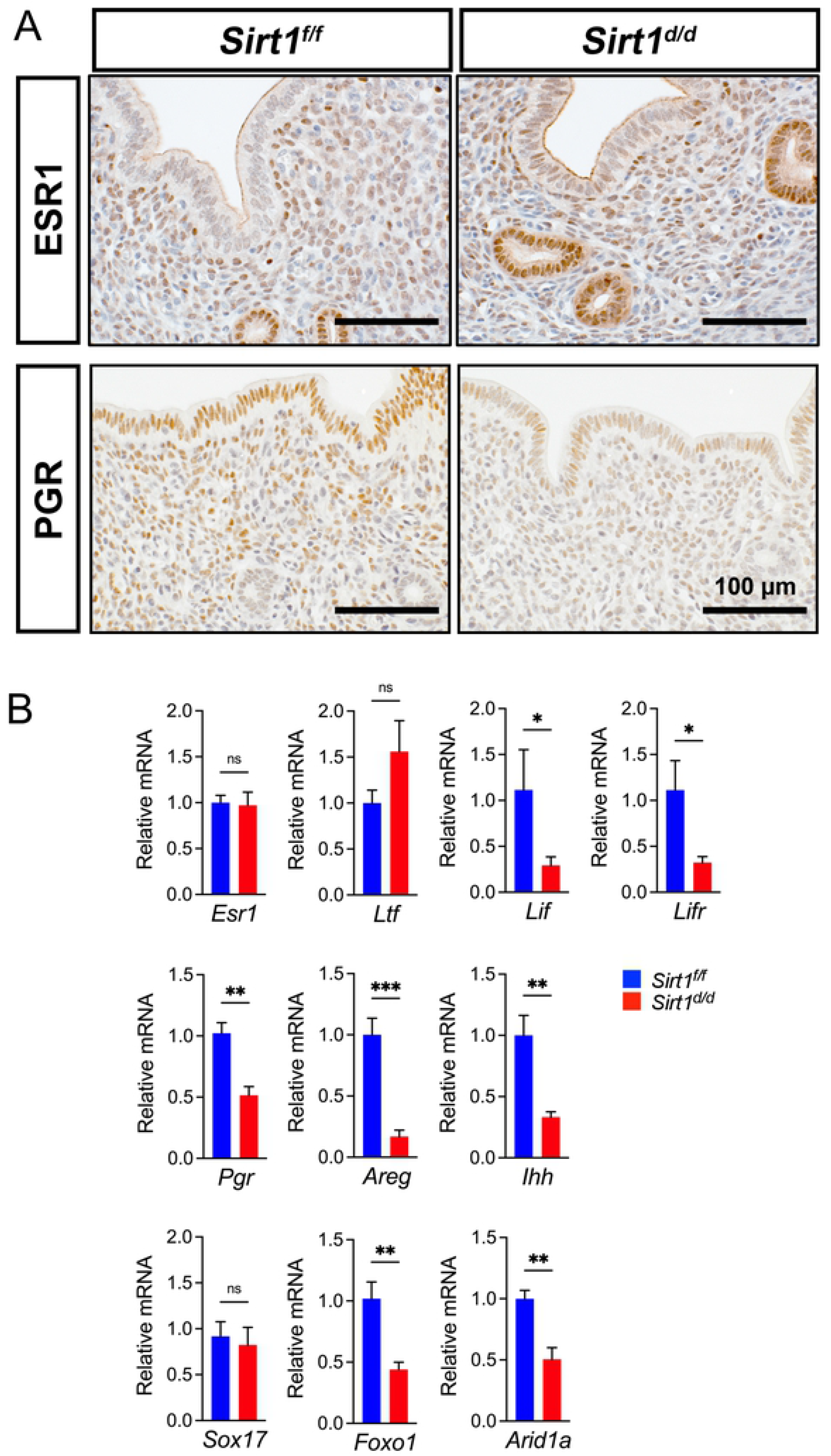
Dysregulated estrogen and progesterone signaling in *Sirt1-*deficient uteri during the window of receptivity to implantation by blastocysts. **(A)** Immunohistochemical staining of ESR1 and PGR in GD 3.5 uteri from *Sirt1^f/f^* and *Sirt1^d/d^* female mice (n=3). **(B)** Quantification of ESR target genes (*Esr1*, *Ltf, Lif, Lifr*), PGR target genes (*Pgr, Areg, Ihh*), and PGR co- regulator genes (*Sox17, Foxo1, Arid1a*) in GD 3.5 uteri from *Sirt1^f/f^* and *Sirt1^d/d^* female mice (n=10). NS, not significant; *, *P*<0.05; **, *P*<0.01; ***, *P*<0.001 (two-tailed *t* test). Data are presented as means ± SEM

### *Sirt1* ablation impairs the Indian hedgehog signaling pathway

Noting that IHH is a major mediator of PGR signaling in the mouse uterus and significantly reduced in SIRT1-ablated uteri, we further investigated effects on downstream of IHH signaling pathways for uterine epithelial-stromal signaling (**Fig 7**). Immunohistochemical analyses of the uteri from 6-8 week old *Sirt1^d/d^* mice at GD 3.5 revealed decreases in the IHH receptors and PTCH1, but not PTCH2 in uterine stromal cells, as well as a reduction in the downstream mediator COUP-TFII (NR2F2) in stratum compactum stromal cells beneath the uterine epithelium (**Fig 7A**). qRT-PCR analyses further confirmed the dysregulation in the IHH signaling pathway, as demonstrated by decreases (*P*<0.05) in expression of mRNAs for *Ptch1, Gli1, Nr2f2* and *Hand2* in GD 3.5 *Sirt1^d/d^* uteri (**Fig 7B**). Consequently, a reduction in decidualization markers *Bmp2* and *Wnt4* were detected (*P*<0.05) in *Sirt1^d/d^* uteri (**Fig 7B**). However, dysregulation of *Fgf* mRNA expression, that accounts for proliferation of epithelial cells, was not consistent in the *Sirt1^d/d^* uteri (i.e., increases in *Fgf1* and *Fgf18*; decreases in *Fgf9*; and no change in *Fgf2, Fgf7* and *Fgf12*) (**Fig 7B**). To determine whether SIRT1 deficiency impacts proliferation of epithelial and stromal cells, expression of proliferation marker Ki67 was determined (**Fig 7B-D**). Based on Ki67 staining of uteri from GD 3.5, *Sirt1^f/f^* mice exhibited minimal levels of cell proliferation (H-score; 1.41+0.85), whereas staining of uterine epithelial from *Sirt1^d/d^* mice was sporadic (H-score; 7.95+2.17; *P*<0.05) (**Fig 7C and D**). It is noteworthy that inhibition of proliferation of uterine epithelial cells does not prevent implantation of blastocysts at GD 5.5 in *Sirt1^d/d^* uteri (**Fig 4A**). Intriguingly, the SIRT1 deficiency also decreases Ki67 expression in the stromal cells (**Fig 7C**) in which expression of *MKi67* is also reduced in the *Sirt1^d/d^* uteri (**Fig 7B**).

**Fig 7.**
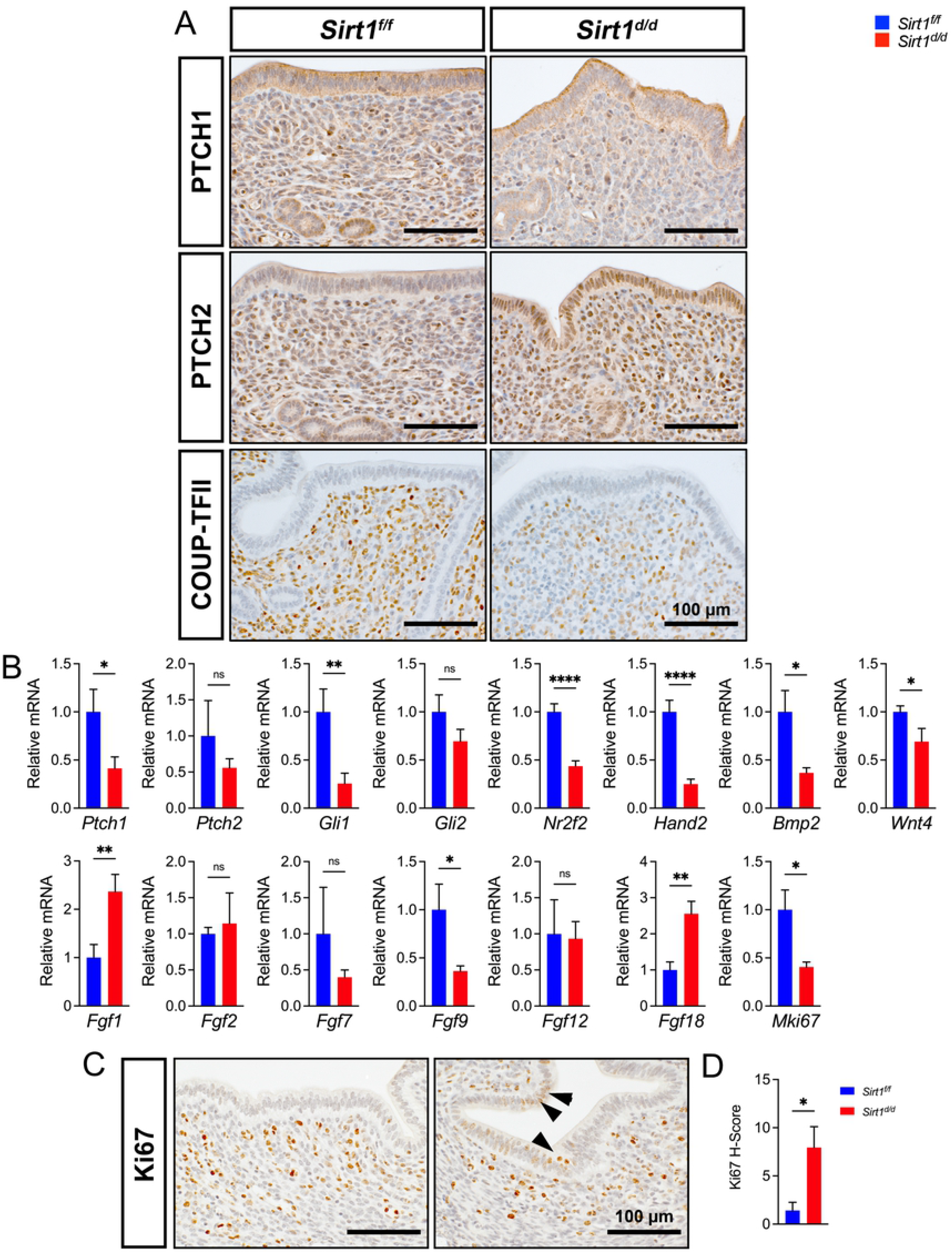
Alteration of Indian Hedgehog (IHH) signaling pathways in *Sirt1-*deficient uterus during the window of receptivity to implantation by blastocysts. **(A)** Immunohistochemical staining of IHH receptors PTCH1, and PTCH2, as well as downstream mediator COUP-TFII in GD 3.5 uteri from *Sirt1^f/f^* and *Sirt1^d/d^* females (n=3). **(B)** Quantification of genes associated with IHH signaling (*Ptch1, Ptch2, Gli1, Gli2, Nr2f2, Hand2*) to prime stromal cells for decidualization (*Bmp2, Wnt4*) and inhibit epithelial proliferation (*Fgf1, Fgf2, Fgf7, Fgf9, Fgf12, Fgf18, Mki67*) in GD 3.5 uteri from *Sirt1^f/f^* and *Sirt1^d/d^* females (n=10). **(C)** Immunohistochemical staining of Ki67 in GD3.5 uteri from *Sirt1^f/f^* and *Sirt1^d/d^* females (n=3). **(D)** H-score quantification of Ki67 in the endometrial section of *Sirt1^f/f^* and *Sirt1^d/d^* females (n=3) at GD 3.5. NS, not significant; *, *P*<0.05; **, *P*<0.01; ****, *P*<0.0001 (two-tailed *t* test). Data are presented as means ± SEM

### SIRT1 regulates the uterine transcriptome to prime the stromal cells for decidualization

To further identify underlying molecular mechanisms of uterine responses during the window of receptivity for blastocyst implantation, we mated *Sirt1^f/f^* and *Sirt1^d/d^* females with intact wildtype males and dissected equivalent pieces of the uterine horns at GD 3.5 for transcriptomic analyses. A total of 6,185 genes were dysregulated at GD 3.5 with 2,577 genes being upregulated and 3,608 genes being downregulated (**S3 Fig A and S1 Table**). Among dysregulated genes by SIRT1 deficiency, several gene clusters were directly associated with uterine receptivity and decidualization as shown in the hierarchical clustering heatmaps, including E2-responsive genes, P4-responsive genes, and hedgehog signaling-associated genes (**Fig 8A-C**). Clustering of E2- responsive genes revealed that expression of *Hgf, Met, Igf1, Vim* and *Mki67* mRNAs was downregulated in uteri of *Sirt1^d/d^* females on GD 3.5, while expression of *Esr1, Esr2, Ltf, Slc27a2, Muc1* and E-Cadherin (*Cdh1*) mRNAs were upregulated (**Fig 8A**). Notably, the majority of P4- responsive genes were downregulated on GD 3.5 in *Sirt1^d/d^* uteri. Those genes were associated with adhesion and invasion of cells (*Itga5, Itga7, Itga8, Itgb1, Itgb2, Itgb3, Mmp2, Mmp7, Mmp11*), and proliferation and differentiation of stromal cells (*Bmp2, Depp1, Gdf10, Hoxa3, Hoxa5, Hoxa9, Hoxa10, Hoxa11, Mcm2, Notch3, Ptger3, Ptgdr2, Sfrp1, Sfrp2, Sfrp4, Tgfb1,Tgfb2, Tgfb3*) (**Fig 8B**). The downregulation of expression of genes in stromal cells provides further evidence for a major defect in mechanisms required for priming stromal cells for decidualization. As expected, IHH pathway associated genes (*Dhh, Gli1, Gli2, Glis1, Glis2, Glis3, Ptch1, Ptch2, Smo, Nr2f2, Hand2* and *Klf15*) were downregulated in *Sirt1^d/d^* uteri on GD 3.5 (**Fig 8C**). Intriguingly, further hierarchical clustering analyses revealed major downregulation of histone proteins and epigenetic modifiers in *Sirt1^d/d^* uteri on GD 3.5, such as *Ezh2, Dnmt1, Dnmt3b, Dnmt3l, Hdac1, Hat1, Hells, Mbd1, Tet1, Tet3*, and *Uhrf1* (**Fig 8D**). Other clusters of genes such as those related to adrenomedullin signaling (*Adm, Ramp1, Ramp2, Calcrl, Ackr3*) (**S3 Fig B**) and hyaluronic acid synthesis (*Has1, Has2, Has3, Hyal2, Cd44*) (**S3 Fig C**) were also downregulated in *Sirt1^d/d^* uteri on GD 3.5; while genes associated with junctional complexes between cells (*Cldn23, Cldn3, Add1, Add2, Add3*) were upregulated (**S3 Fig D**).

**Fig 8.**
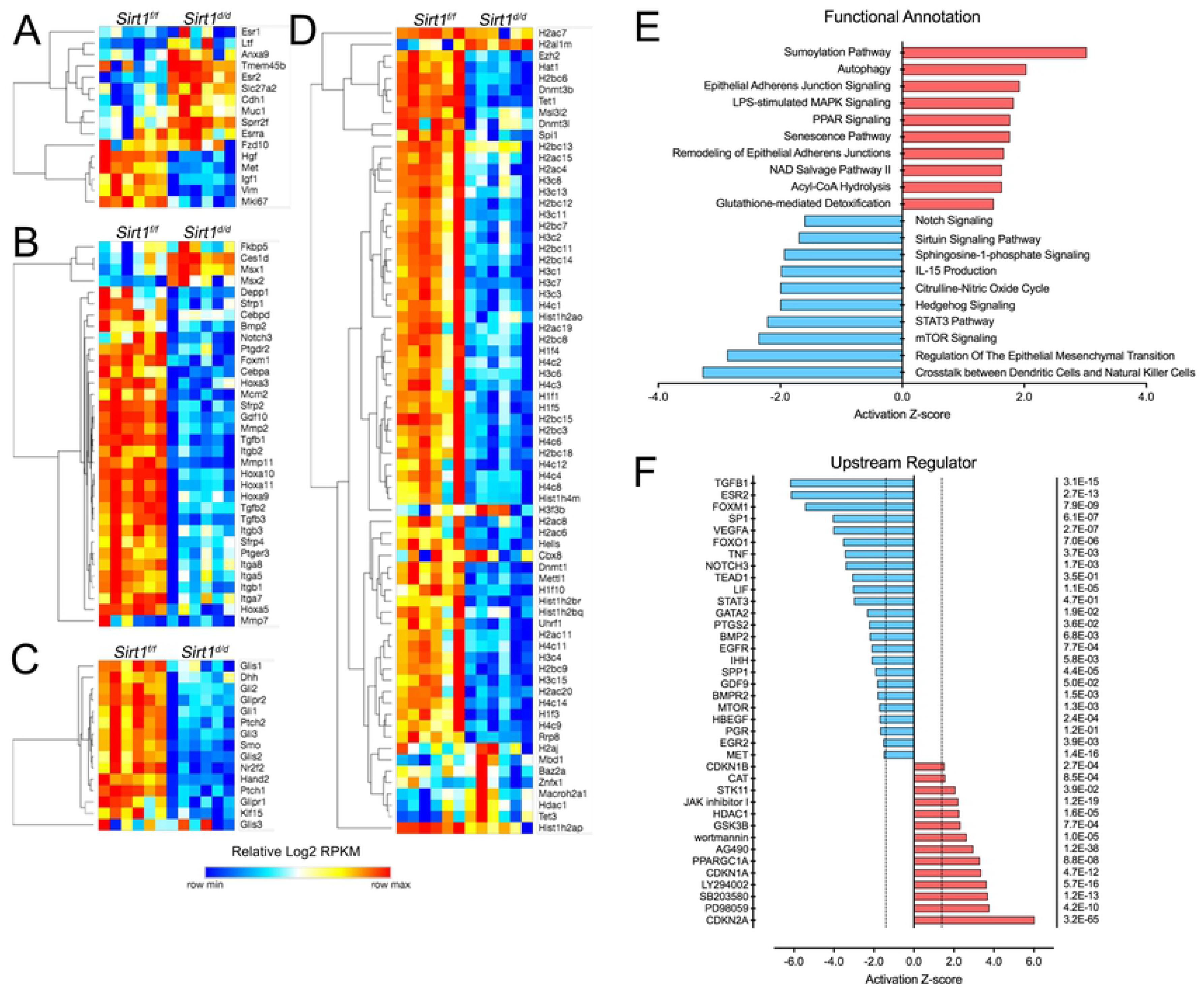
SIRT1 regulation of the uterine transcriptome. **(A-D)** Hierarchical clustering heatmaps depicting the expression profiles of E2-responsive genes (**A**), P4-responsive genes (**B**), IHH signaling pathway (**C**), as well as histone proteins and epigenetic modifiers (**D**) that are differentially expressed between *Sirt1^f/f^* and *Sirt1^d/d^* mouse uteri at GD 3.5 generated from RNA-seq analysis (n=6). **(E)** Enrichment of canonical pathways in differentially expressed genes between *Sirt1^f/f^* and *Sirt1^d/d^* mouse uteri at GD 3.5 by IPA analysis (n=6). **(F)** Enrichment of upstream transcriptional regulators for SIRT1-deficient transcriptome at GD 3.5 by IPA analysis (n=6). Activation Z-score denotes the direction of change in which Z-score > 1.5 indicates activation, and Z-score < 1.5 indicates inhibition (*P*<0.05). Also see **S3 Fig**.

To further identify changes in biological functions of uteri in mice with a SIRT1-deficient transcriptome and obtain evidence for the bases for regulation of expression of particular clusters of genes, Ingenuity Pathway Functional Annotation and Upstream Regulator analyses were performed. The top-enriched functional annotations for the aberrantly controlled gene list (6,185 genes) included, but were not limited to activation of SUMOylation, autophagy, senescence and NAD salvage pathways; as well as inhibition of crosstalk between dendritic cells and natural killer cells, the epithelial-mesenchymal transition, hedgehog signaling, Sirtuin signaling and Notch signaling pathways (**Fig 8E**). Meanwhile, the top-enriched upstream regulators of the SIRT1- deficient transcriptome included, but were not limited to: (1) inhibition of TGFB1, ESR2, FOXO1, NOTCH3, LIF, STAT3, GATA2, PTGS2, BMP2, EGFR, IHH, SPP1, GDF9, HBEGF and PGR; and (2) activation of CDKN2A, PD98059 (MEK1 inhibitor), SB203580 (p38 MAPK inhibitor), and LY294002 (PI3K inhibitor). The entire list of functional annotations and upstream regulators is provided in **S2 and S3 Tables**.

### Uterine-specific deletion of *Sirt1* results in premature uterine aging

Next, we performed a correlation search to determine which available dataset in the Illumina BaseSpace Correlation Engine database best characterized the gene expression profile of *Sirt1^d/d^* uteri during the window of receptivity to implantation of blastocysts. The top-ranking datasets sharing similar gene expression patterns with the SIRT1-regulated transcriptome at GD 3.5 were uterine transcriptomes of (1) advanced maternal age; (2) *Foxa2* knockout, KO; (3) *Foxo1* KO; (4) *Foxl2* overexpression, OE; (5) *Wnt4* KO; (6) *Sox17* KO; (7) *Bmp2* KO; (8) *Egfr* KO; (9) *Nr2f2* KO; (10) *Esr1* KO; and (11) *Pgr* KO (**Fig 9A**). With the strongest positive correlation, the *Sirt1^d/d^* and aged transcriptomes share 1,157 deregulated genes with 455 genes upregulated and 639 genes downregulated (**Fig 9B and S4 Table**). The overlaps of commonly deregulated genes between *Sirt1* KO and other top-ranking uterine transcriptomes are listed in **S4 Fig**. These findings support our hypothesis that SIRT1 plays an essential role in governing age-related changes in uterine receptivity for implantation of blastocysts and stromal cell decidualization, and that uterine-specific ablation of SIRT1 accelerates premature aging of the uterus in mice. In addition, analysis of the deposition of age-related fibrillar collagens, as measured by PSR staining, showed that at GD 3.5 *Sirt1^f/f^* mice exhibited minimal staining of fibrillar type I and III collagens in the uterine stroma. On the other hand, both young *Sirt1^d/d^* and aged wildtype uterine stroma stained positive for type I and III collagens and thus exhibited age-related uterine stromal fibrosis, suggesting that a SIRT1 deficiency accelerates premature uterine aging.

**Fig 9.**
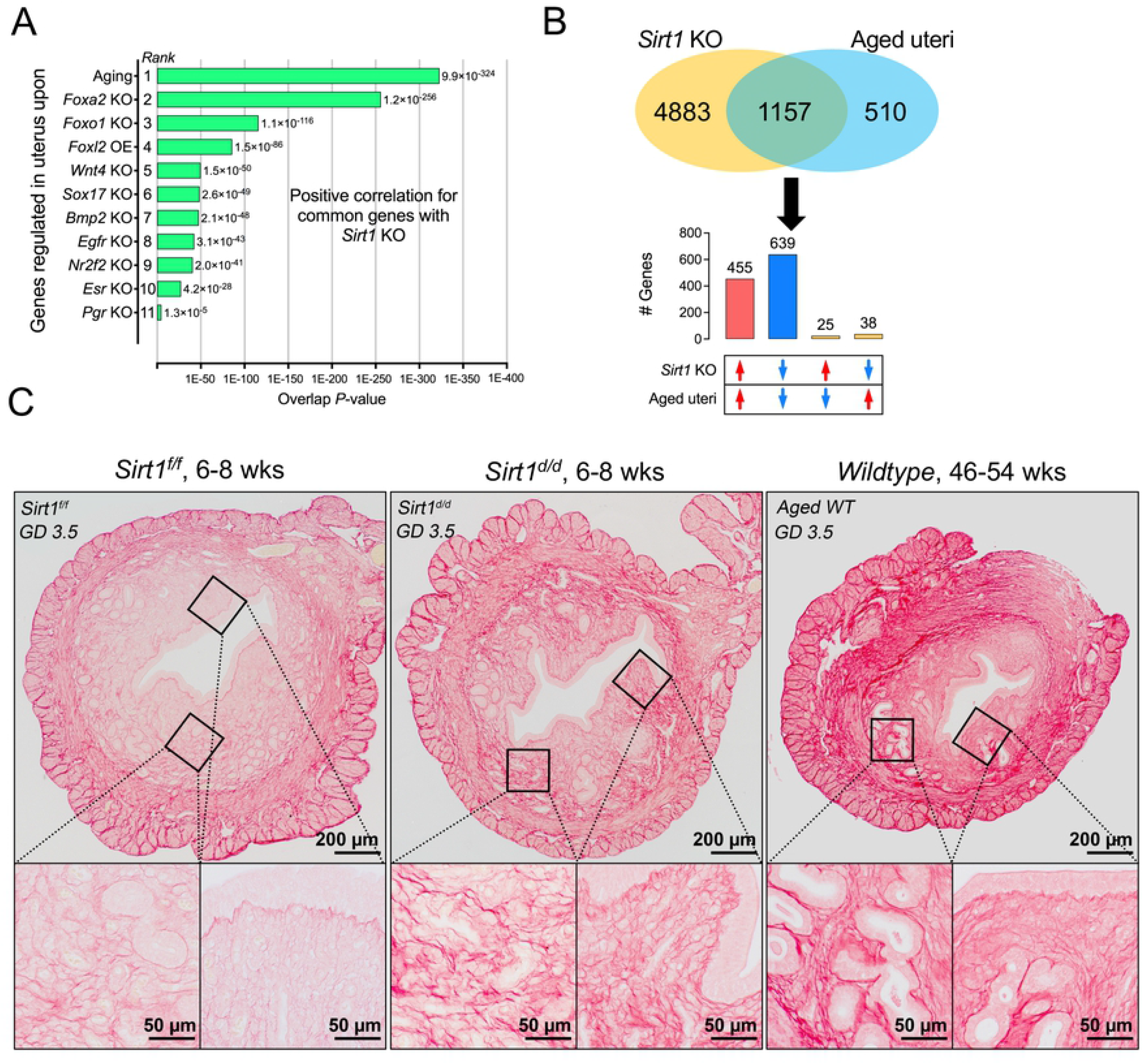
Uterine-specific deletion of *Sirt1* results in premature uterine aging. **(A)** Top uterine transcriptomic biosets overlapping with *Sirt1* KO transcriptome generated by NextBio gene expression profile comparison. **(B)** Overlaps and correlations of *Sirt1* KO with aged transcriptomes at GD 3.5, identifying 1,094 genes commonly deregulated in *Sirt1* KO and aged uteri. Also see **S4 Fig.** for a separate comparison between SIRT1 KO and other top similar transcriptomes. **(C)** Hierarchical clustering heatmap of commonly deregulated genes among top uterine transcriptomic biosets based on the 1,094 genes depicted in (B). **(D)** Picrosirius Red staining of uteri from young *Sirt1^f/f^*, young *Sirt1^d/d^* and aged wildtype female mice, depicting accelerated disposition of aging-related fibrillar type I and III collagens in SIRT1-deficient uterine stroma. Inserts indicate high magnification of box area. KO, knockout; OE, overexpression.

## Discussion

Advanced maternal age negatively impacts reproductive outcomes, leading to pregnancy complications such as stillbirth, preterm birth or IUGR, as well as a range of developmental defects to the newborn. These adverse pregnancy outcomes are becoming more widespread as women worldwide are delaying childbearing for reasons ranging from higher educational attainment, effective contraception, and financial and economic concerns [50, 51]. Recent studies with mice have shed light on the decidualizing uterus that has a blunted PGR signaling response as the major cause of age-related reproductive decline unrelated to oocyte quality [17, 52]. It is noteworthy that the number of live offspring declines in older females despite unchanged numbers of early implantation sites, suggesting that embryonic losses must occur post-implantation [53, 54]. Here we demonstrate that uterine SIRT1 is a critical driver of age-related PGR action required for uterine epithelial-stromal crosstalk, thereby regulating spacing and implantation of blastocysts, as well as uterine stromal decidualization and placentation (**Fig 10**).

**Fig 10.**
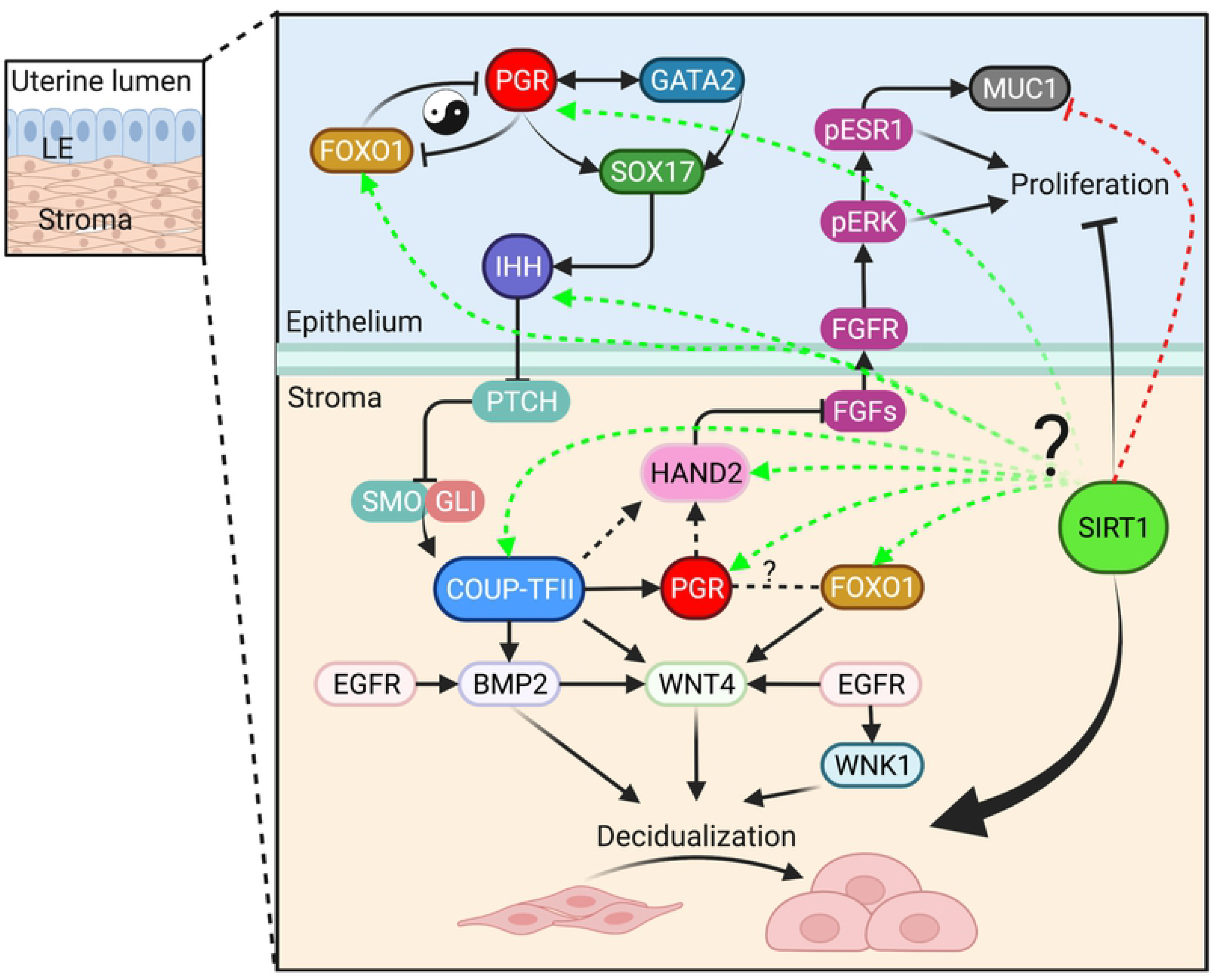
SIRT1 regulates uterine epithelial-stromal interactions to fine-tune epithelial proliferation and direct stromal cell decidualization.

SIRT1 is a pivotal regulator of chromatin structure [18], senescence [22, 24] and inflammation [29, 30]. Alterations in SIRT1 signaling is associated with aging in various tissues outside of the uterus [22–28] as SIRT1 levels are lower in cells that have been serially spilt or in dividing tissues of aged mice, such as thymus and testis [55]. Recent studies have shown that oocyte-specific ablation of SIRT1 induces premature aging by compromising oocyte quality [56], whereas overexpression of SIRT1 in the oocytes extends ovarian lifespan in aged mice [57]. Here we determine that aged uteri exhibit intrinsic defects in the temporal progression of decidual differentiation, accompanied by a decline in *Sirt1* mRNA levels. During the peri-implantation period of pregnancy in young mice, *Sirt1* mRNAs were expressed in the epithelial and stromal compartment of the endometrium. In addition, we provide novel *in vivo* evidence indicating that SIRT1 is the common target positively regulated by SOX17 and FOXO1, two key co-regulators of PGR for implantation and decidualization [39, 40]. These results led us to hypothesize that SIRT1 plays a critical role in governing age-related PGR actions required for implantation and decidualization.

To test this hypothesis, we generated mice with uterine-specific ablation of Sirt1, which are subfertile with low litter size and frequency of producing offspring, as well as delivering more stillborn pups. The fact that the majority of *Sirt1^d/d^* mothers become sterile at 25 weeks of age after giving birth to the 3^rd^ litter document that they experience premature reproductive aging. The majority of pregnancy complications associated with advanced maternal age, such as IUGR and stillbirth, often share underlying pathogenesis due to deficiencies in placentation, independent of oocyte health [58]. Previous studies have pinpointed the cause to a more profound defect in stromal cell decidualization that is linked to heterogeneity in expression of both PGR and ESR responsive genes at GD 3.5, and the window of receptivity for implantation of blastocysts in the aged uteri [17]. Concordant with these observations, our results revealed that a uterine SIRT1 deficiency impairs stromal cell decidualization, spacing and implantation of blastocysts, deficiencies in uterine decidualization, abnormal placentation and adverse pregnancy outcomes. We then sought to determine changes to the decidualizing uterus that interfere with spacing, implantation, and placentation in SIRT1-deficient mice.

To address this question, we first identified changes at GD 5.5 in SIRT1-deficient mice responsible for the initial progression of stromal cell decidualization. In SIRT1-deficient uteri, the expression pattern of PTGS2 was not altered in DSCs closely surrounding the entire blastocyst, and FOXO1 remained in the intact uterine LE at the fetal-maternal interface, suggesting a delay in implantation. However, there were decreases in PGR in stromal cells proximal to the blastocyst, as well as the restricted area of DSCs indicative of dysregulated decidualization in uteri of SIRT1- deficient mice. In addition, FOXO1 expression was low in uterine LE and S on the mesometrial side of the uteri of *Sirt1^d/d^* mice on GD 5.5, indicating that the SIRT1 deficiency compromised stromal cell decidualization, possibly via FOXO1. Given that FOXO1 regulates SIRT1 and that they are similar in phenotypes [39], interactions between FOXO1 and SIRT1 may account for stromal cell decidualization.

Next, we demonstrate that this delay in implantation and uterine decidualization can be traced back to the very early stages of hormonal priming of the uterus at GD 3.5. Together with our analyses of the transcriptome, the SIRT1 deficiency disrupted ESR signaling and suppressed PGR signaling in the uterus during the window of receptivity for implantation of the blastocyst. Both LIF and IHH signaling were altered in this mouse model to yield a phenotype similar to that for SOX17-deficient uteri [40]. LIF is produced by the uterine GE and via receptors in the uterine LE to regulate implantation of blastocysts and stromal cell decidualization [59]. IHH initiates the epithelial-stromal crosstalk required for cessation of proliferation of uterine epithelial cells, blastocyst implantation and stromal cell decidualization [60]. However, inhibition of proliferation of uterine epithelial cells was not severely compromised because blastocysts were able to implant in *Sirt1^d/d^* uteri. Instead, failure of stromal cell decidualization is evident due to dysregulation of the IHH-COUP-TFII-BMP2-WNT4 signaling pathway involving actions requiring epithelial (ePGR) and stromal (sPGR) PGR. Moreover, adrenomedullin is a highly conserved peptide hormone required for intrauterine spacing of blastocysts during early pregnancy [61, 62]. Hyaluronan (hyaluronic acid; HA)-CD44 interactions promote proliferation and differentiation of stromal cells [63] and its metabolism fine-tunes the avascular niche and maternal vascular morphogenesis in the implantation chamber of blastocysts [64]. Thus, downregulation of the ADM and HA signaling pathway in SIRT1-deficient uteri may also contribute to defects in blastocyst spacing and implantation, as well as stromal cell decidualization. Downregulation of a significant number of histone proteins and dysregulation of epigenetic modifiers further suggests that the uterine SIRT1 deletion triggers DNA damage and perturbations due to epigenetic modifications, respectively.

Integrating our results with published results from studies of the model for uterine aging effects and key regulators on stromal decidualization and/or implantation failure, we found significant overlaps with uterine transcriptomes at the same days of pregnancy, as well as master regulators of decidualization, notably those for the PGR signaling pathway including *Foxo1, Foxl2, Wnt4, Sox17, Bmp2, Egfr and Nr2f2*. Interestingly, the uterine FOXA2-deficient transcriptome ranked second as significantly overlapping with the uterine SIRT1-deficient transcriptome. Given that SIRT1 deletion did not affect FOXA2 expression, but downregulates LIF, it is possible that SIRT1 is a key target downstream of FOXA2, the critical regulator of uterine glandular function required for implantation and stromal cell decidualization [65]. The significant overlaps with uterine transcriptomes of aged mice suggests that by deleting SIRT1 a genetic aging model partially, if not entirely, recapitulating physiological aging in the uterus was produced. Furthermore, accelerated disposition of aging-related fibrillar type I and III collagens in *Sirt1*-deficient uteri substantiate the functional role of SIRT1 in influencing the adaptability of uterine functions to support pregnancy during reproductive aging.

Collectively, results of our study highlight the importance of uterine SIRT1 as the indispensable age-related regulator of PGR actions for implantation, decidualization and placentation in mice. PGR is the master regulator for the establishment and maintenance of pregnancy [66]; however, a significant diminution in PGR results in a blunted hormonal response as the uterus ages [17]. The underlying molecular mechanisms that diminish expression of PGR likely account for uterine aging, a mechanism that has remained elusive. By deleting uterine SIRT1, we produced a genetic model for research on premature uterine aging due to blunted PGR responses that are similar to those associated with physiological aging. Although the extent to which these findings from the mouse model translates to humans, it is evident that results of the present study significantly advance understanding of molecular changes responsible for blunting PGR actions in aging uteri, independent of potential changes in oocyte quality. Future studies on how SIRT1 interacts with factors in the PGR signaling pathway (e.g. SIRT1-responsive acetylome, and epigenetic modification) may identify gene signatures in the uterus that are sensitive to reproductive aging. With that new knowledge, research can pursue strategies to counteract adverse effects of aging on outcomes of pregnancy.

## Materials and Methods

### Animals care and use

All animal experiments were conducted in full compliance with the Guide for the Care and Use of Laboratory Animals published by the National Institute of Health and with approval of the Institutional Animal Care and Use Committee (IACUC) at North Carolina State University under protocol 19-040-B. C57BL/6J mice were used throughout this study. “Young” females were generally 6-8 weeks old. Timed matings were set up with standard C57BL/6J males between 8 and 16 weeks of age. The morning when the vaginal plug was detected was designate gestational day (GD) 0.5. “Aged” males were 46-54 weeks old. The uterine-specific *Sox17* knockout mice (*Sox17^d/d^*) were generated by crossing mice carrying a *Sox17^f/f^* allele (stock no: 007686, the Jackson Laboratory, Bar Harbor, ME) with *Pgr^Cre/+^* mice ([43]; a kind gift from Drs. Francesco J. DeMayo at NIEHS, Research Triangle Park, NC, and John P. Lydon at Baylor College of Medicine, Houston, TX). The uterine-specific *Foxo1* knockout mice (*Foxo1^d/d^*) were generated by crossing mice carrying a *Foxo1^f/f^* allele (stock no: 024756, the Jackson Laboratory Bar Harbor, ME) with *Pgr^Cre/+^* mice. The uterine-specific *Sirt1* knockout mice (*Sirt1^d/d^*) were generated by crossing mice carrying a *Sirt1* exon 4 floxed allele (*Sirt1^f/f^*; [41, 42]; obtained from Dr. Xiaoling Li at NIEHS, Research Triangle Park, NC.) with *Pgr^Cre/+^* mice.

### Animal fertility assay

Conditional knockout and control littermate females at approximately six weeks of age were housed individually and continuously with wildtype C57BL/6J males. Mating was confirmed by the presence of vaginal plugs. Fertility was assessed by monitoring the frequency of litters and litter sizes for a six-month period.

### Artificial decidualization

To assess the ability of the stroma to undergo differentiation and proliferation independent of implantation by a blastocyst, (1) aged or young female mice; as well as (2) *Sirt1^f/f^* or *Sirt1^d/d^* female mice were ovariectomized and treated with exogenous hormones to mimic pregnancy before applying a manual stimulus to a single uterine horn (protocol outlined previously by Finn and Martin, 1972)[67]. After ovariectomy and 2 weeks of rest to eliminate endogenous ovarian steroids, mice were administered E2 (E8875, Sigma-Aldrich, St. Louis, MO; 100 ng per mouse) as daily injections for 3 days. After 2 days of rest, mice were treated with E2 (6.7 ng per mouse) and P4 (P0130, Sigma-Aldrich; 1 mg per mouse) for 3 days. On the third day (DD0), mice were administered a single injection of 0.05 ml sesame oil into the right uterine horn. Mice were administered E2 and P4 for 5 more days and killed on the fifth day (DD5). Uterine wet weights for the stimulated and control horns were recorded. Weight ratios were calculated by dividing stimulated horn weight by unstimulated horn weight.

### Recovery of blastocysts

Adult *Sirt1^f/f^* or *Sirt1^d/d^* females were mated overnight with wildtype males. Females were separated from males the following morning. The presence of a vaginal plug was considered 0.5 day post coitum (GD 0.5). Uterine horns were harvested at GD 3.5. Blastocysts were recovered by flushing the uterine lumen with phosphate-buffered saline (PBS) and the numbers were counted under a bright-field microscope.

### Blastocyst implantation, spacing, invasion and pseudopregnancy

The ability of blastocysts to undergo implantation was determined by mating 8-week-old females with wildtype males. The morning a vaginal plug was observed was considered 0.5 days post coitum (GD 0.5). Mice were sacrificed at GD 5.5, 6.5 and 9.5. Uteri were excised, imaged, and decidua tissue weights at GD 6.5 were determined. Spacing and invasion of blastocysts were calculated using Image J (NIH) and the quantitative method depicted in Figure 4 C and G, respectively. Briefly, the distance between the middle of each of two adjacent implantation sites was measured to generate a standard deviation (SD) for each uterus. High SD indicates uneven spacing. For assessment of invasion of blastocysts, the distance from the blastocyst to the myometrium on the mesometrial side (a) or anti-mesometrial side (b) were measured to generate a distance ratio (a/b). A high distance ratio indicates significant invasion by the blastocysts. Uteri were fixed in 4% v/v electron microscopy grade paraformaldehyde (PFA) in PBS for histology. Analysis of uterine gene expression during pseudopregnancy was accomplished by mating 8-week-old female mice with vasectomized male mice. The morning of detection of the postcoital vaginal plug was designated pseudopregnant day (PPD) 0.5. The mice were sacrificed on PPD 4.5.

### RNA *in situ* hybridization analyses

RNAscope *in situ* hybridization (ISH; Advanced Cell Diagnostic, Newark, CA) was performed according to the manufacturer’s instructions using PFA-fixed uterine tissues (∼5 μm) and Mm-SIRT1 probe (418341) for murine *Sirt1* mRNA. A bacterial DapB gene probe (310043) was used as the negative control. Following hybridization, slides were washed and probe binding visualized using the HD 2.5 Red Detection Kit (322360-USM). Sections were briefly counterstained with hematoxylin before being dehydrated and affixing coverslips with Permount.

### RNA isolation

Frozen tissue was homogenized in 1 ml of TRIzol reagent (Thermo Fisher) using a Bead Mill 24 homogenizer (Thermo Fisher), two times at 4.5 m/s for 30 s, and rested intermittently for 20 s on ice. Homogenates were centrifuged for 10 min at 12,500 *g* at 4°C to pellet cellular debris. The supernatant was transferred to a 1.5-ml centrifuge tube and mixed with 200 μl of 1-bromo-3-chloropropane by manually shaking for 20 sec. The tube was then incubated at room temperature for 3 min, and centrifuged for 18 min at 12,500 *g* at 4°C. The upper aqueous phase (∼400 μl) was carefully removed, placed into a new 1.5-ml tube, and mixed with 200 μl of chloroform by shaking the tubes for 20 sec. Samples were rested for 3 min at room temperature and subsequently centrifuged at 21,000 *g* for 18 min at 4°C. Approximately 500 μl of the aqueous layer was transferred to a new tube and mixed with equal parts of 70% ethanol. This mix was filtered in columns from the RNeasy Mini kit (Qiagen, Valenica, CA). Columns were washed once with 700 μl of RW1 buffer (1053394; Qiagen) and three times with 500 μl of RPE buffer (1018013; Qiagen). The RNA was eluted with 30 μl RNase-free water. The quantity and quality of total RNA were determined using spectrometry and denaturing agarose gel electrophoresis, respectively.

### Quantitative real-time PCR analyses

RNA was reverse transcribed into cDNA using the Moloney Murine Leukemia Virus (M-MLV; Thermo Fisher) according to the manufacturer’s instructions. Quantitative RT-PCR (qRT-PCR) was performed using the CFX Connect Real-Time PCR Detection System (Bio-Rad, Hercules, CA, USA) and 1) the SsoAdvanced™ Universal SYBR® Green Supermix (1725274; Bio-Rad) with oligonucleotide primers synthesized by Integrated DNA Technologies (IDT; Coralville, IA, USA), or 2) the SsoAdvanced™ Universal Probes Supermix (1725284; Bio-Rad) with Taqman probes (Applied Biosystems). Delta delta Ct values were calculated using 18 S and ACTB control amplification results to determine relative variations in expression of mRNAs per sample. Information for all primers and Taqman probes is provided in Table **S5**.

### Western blot analyses

Extracted proteins (30 μg/sample) were denatured, separated using SDS-PAGE (4 to 12 % gradient gel at 150 V for 2.5-3 h) and transferred to a nitrocellulose membrane overnight (∼16 h) at 20 V using the Bio-Rad Transblot (Bio-Rad). Membranes were blocked in 5 % fat-free milk in 20 mmol/l Tris, 150 mmol/l NaCl, pH 7.5, and 0.1 % Tween-20 (TBST) for 3 h and then incubated with a primary antibody for SIRT1 (1:1000; #2028) or ACTB (1:5000; #4970) at 4°C overnight with gentle rocking. After washing three times (10 min per time) with TBST, the membranes were incubated for 2 h with a secondary antibody (horseradish-peroxidase-linked anti-rabbit IgG) at 1:10 000 dilutions. The membranes were then washed with TBST, followed by development using enhanced chemiluminescence detection (SuperSignal West Pico) according to the manufacturer’s instructions. Western blots were analyzed by measuring the intensity of light emitted from correctly sized bands under ultraviolet light using a ChemiDocTM MP Imaging System (Bio-Rad). Multiple exposures of each Western blot were performed to ensure the linearity of chemiluminescence signals.

### Histological and immunohistochemical staining

At the time of sacrifice, a mid-portion of the uterine horn was fixed in 4% v/v PFA and embedded in paraffin wax. Embedded tissues were sectioned at 5 μm and baked 1 h at 60°C. Upon cooling, slides were dewaxed using Citrisolv clearing agent (22-143-975, Thermo Fisher) in a decreasing gradient of pure ethanol. For hematoxylin and eosin (H&E) staining, tissues were adequately stained with H&E and were then dehydrated before coverslips were applied. For immunohistochemistry, antigen retrieval was performed according to the manufacturer’s instructions (Vector Labs Antigen Unmasking Solution H-3300). Endogenous peroxide was blocked using 3% hydrogen peroxide diluted in methanol. The tissue was blocked with 5-10% normal donkey serum before application of the primary antibody overnight at 4°C. Secondary antibody was diluted in 1% bovine serum albumin at a concentration of 1:200 when required. The ABC reagent was applied to tissues according manufacturer’s instructions (Vector Labs ABC PK-6100). Signal was developed using Vector Labs DAB ImmPACT staining according to manufacturer’s instructions (Vector Labs SK-4105). Tissue was counterstained with hematoxylin and dehydrated before affixing coverslips. A semiquantitative grading system (H-score) was used to compare the immunohistochemical staining intensities [40, 68]. The H-score was calculated using the following equation: H-score = Σ Pi (i), where i = intensity of staining with a value of 1, 2, or 3 (weak, moderate or strong, respectively) and Pi is the percentage of stained cells for each intensity, varying from 0 to 100%. Information for all antibodies is provided in Table **S6**. For Picosirius red (PSR) staining, tissue sections were deparaffinized in Citrisolv and then rehydrated in a series of graded ethanol baths (100%, 70% and 30%). Slides were washed in water and immersed in a PSR staining solution prepared by dissolving Sirius Red F3BA (Direct Red 80, C.I. 357.82, Sigma-Aldrich) in a saturated aqueous solution of picric acid (P6744, Sigma-Aldrich) at 0.1% w/v. After 40 min of incubation at room temperature, the slides were then incubated in 0.05 M hydrochloric acid for 90 sec. Slides were washed in water, dehydrated in 100% ethanol and cleared in Citrisolv for 15 min each before affixing coverslips with Permount.

### RNA-seq and bioinformatic analyses

For RNA-seq from uteri at GD 3.5, total RNA (350 ng) was prepared (see **RNA isolation**) to generate a library using the TruSeq Stranded Total RNA with Ribo-Zero Plus kit (Illumina) following the manufacturer’s instructions. Indexed libraries were sequenced with a 75 bp single-end protocol on an Illumina NextSeq500 sequencer. Raw fastq data were mapped to the *Mus musculus* GRCm38 genome assembly using HISAT2 v2.2.0. Data were quantitated at a protein-coding mRNA level using Partek Flow bioinformatics software (https://www.partek.com/partek-flow/), and normalized according to total read count (reads per kilobase of transcript per million mapped reads, RPKM), and 75% distribution. Differential expression was calculated using DESeq2. Transcripts with the average RPKM >1 in at least 1 group, *q* < 0.05, and at least 1.5-fold difference in RPKM were defined as differentially expressed genes (DEGs). Files were deposited to GEO Datasets under accession number GSE186065.

### Data analyses

The DEGs identified using RNA-seq were analyzed using Ingenuity Pathway Analysis software (IPA, http://www.ingenuity.com) and Database for Annotation, Visualization, and Integrated Discovery (DAVID, http://david.ncifcrf.gov/). Similar transcriptomes to the *Sirt1^d/d^* gene expression profile were identified by searching available datasets in the NextBio database (https://www.nextbio.com/). The ChIP-Seq data were analyzed using the Cistrome software (http://cistrome.org/ap/) and visualized on UCSC Genome Browser (https://genome.ucsc.edu).

The GraphPad Prism software (v9.2.0) was implemented for one-way ANOVA, multiple comparison test, two-tailed *t* test and χ^2^ test analyses. Hierarchal clustering heatmaps were generated using Partek Genomics Suite 6.6 software and MORPHEUS (https://software.boardinstitute.org/morpheus).

## Author Contributions

Conceptualization: MJC, XW.

Data curation: MJC, HY, SP.

Formal analysis: MJC, GH, XW.

Investigation: MJC, HY, SP, XW.

Writing – original draft: MJC, XW.

Writing – review & editing: HY, GH, XL, MH, XW

## Acknowledgements

Contributions of the graduate students and postdoctoral fellows from the Laboratory of Reproductive and Developmental Biology are gratefully acknowledged. The authors also thank Drs. Francesco J. DeMayo and Fuller W Bazer for scientific discussion, and Drs. Francesco J. DeMayo and John P. Lydon for providing *Pgr*-Cre mice.

---

**S1 Fig.**
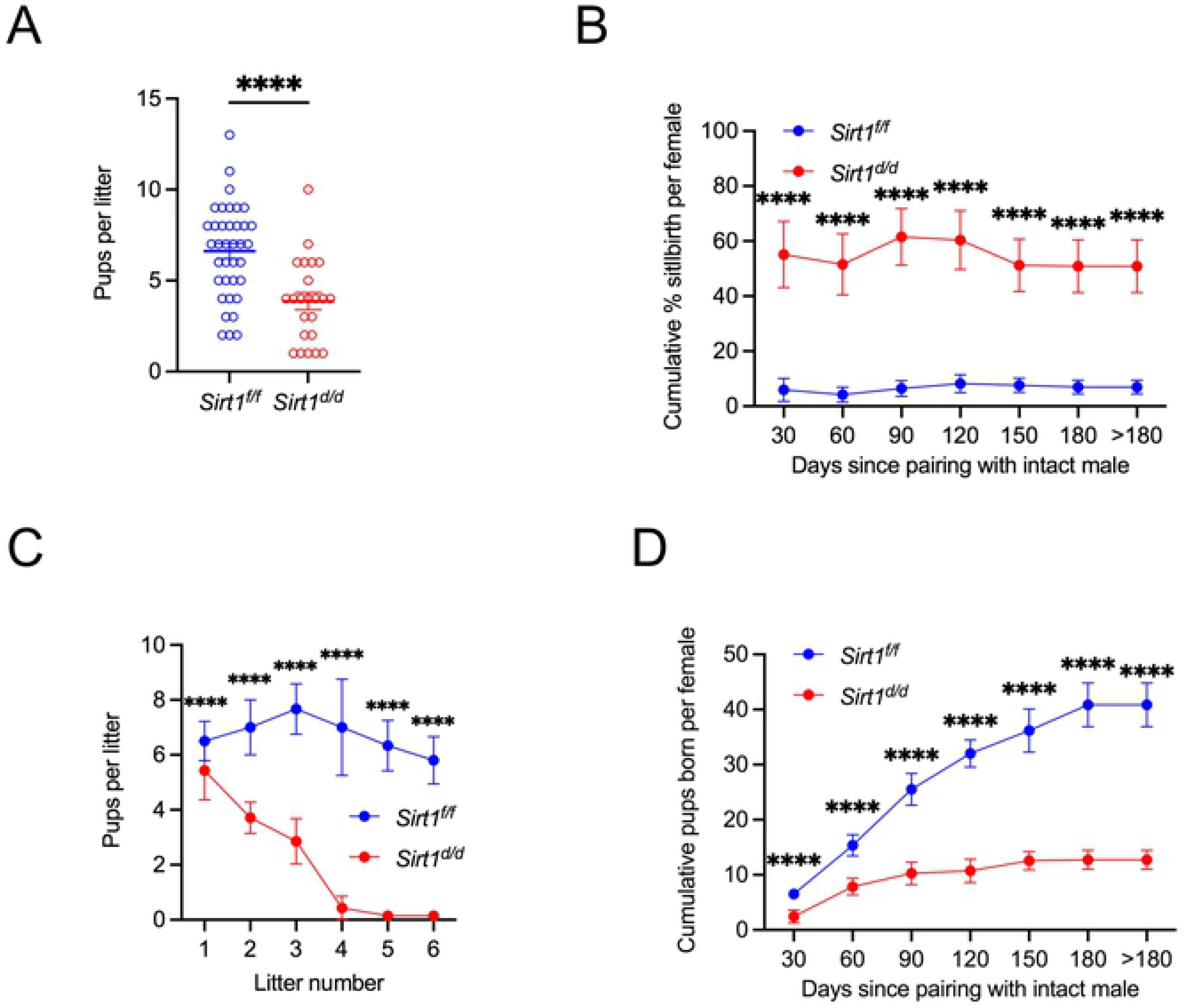
Impact of Sirt1 KO on the number of total and stillborn pups. **(A)** Total pups born per litter. **(B)** Cumulative numbers of stillborn pups since pairing. The mean number of stillborn pups produced up to each time point is denoted along with the respective error bar, with >180 d representing the total number of stillborn pups produced. (**C**) Total pups born in a specific number of litters. **(D)** Cumulative total pups born since pairing. The mean number of pups produced up to each time point is denoted by each error bar, with >180 d representing the total number of pups produced. n=6 of *Sirt1^f/f^*; and n=7 of *Sirt1^d/d^*. ****, *P*<0.0001 (Two-way ANOVA with Tukey’s multiple comparisons test) at specific time points for contrast between *Sirt1^f/f^* (in blue, n=6) and *Sirt1^d/d^* (in red, n=7) female mice. Data are presented as means ± SEM

**S2 Fig.**
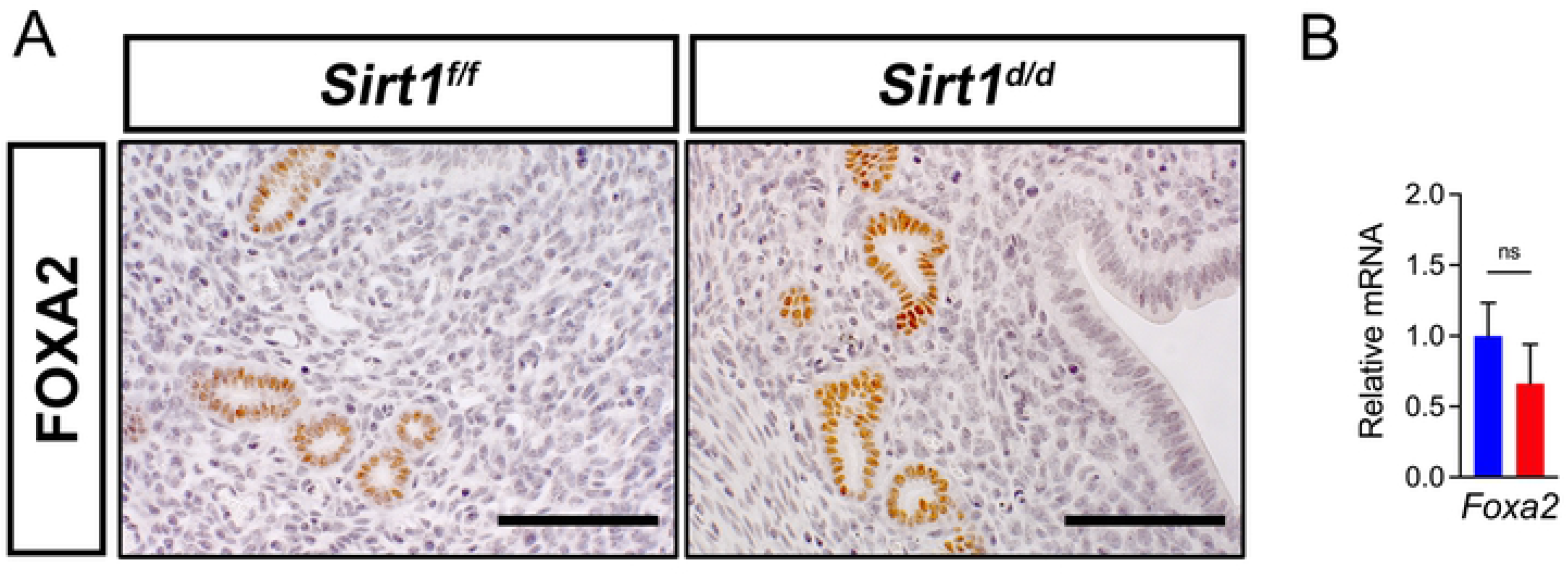
Immunohistochemical (A) and qRT-PCR analyses (B) of *Foxa2* gene in uteri from *Sirt1^f/f^* and *Sirt1^d/d^* female mice on GD 3.5. Scale bar: 100μm. NS, not significant (two-tailed *t* test).

**S3 Fig.**
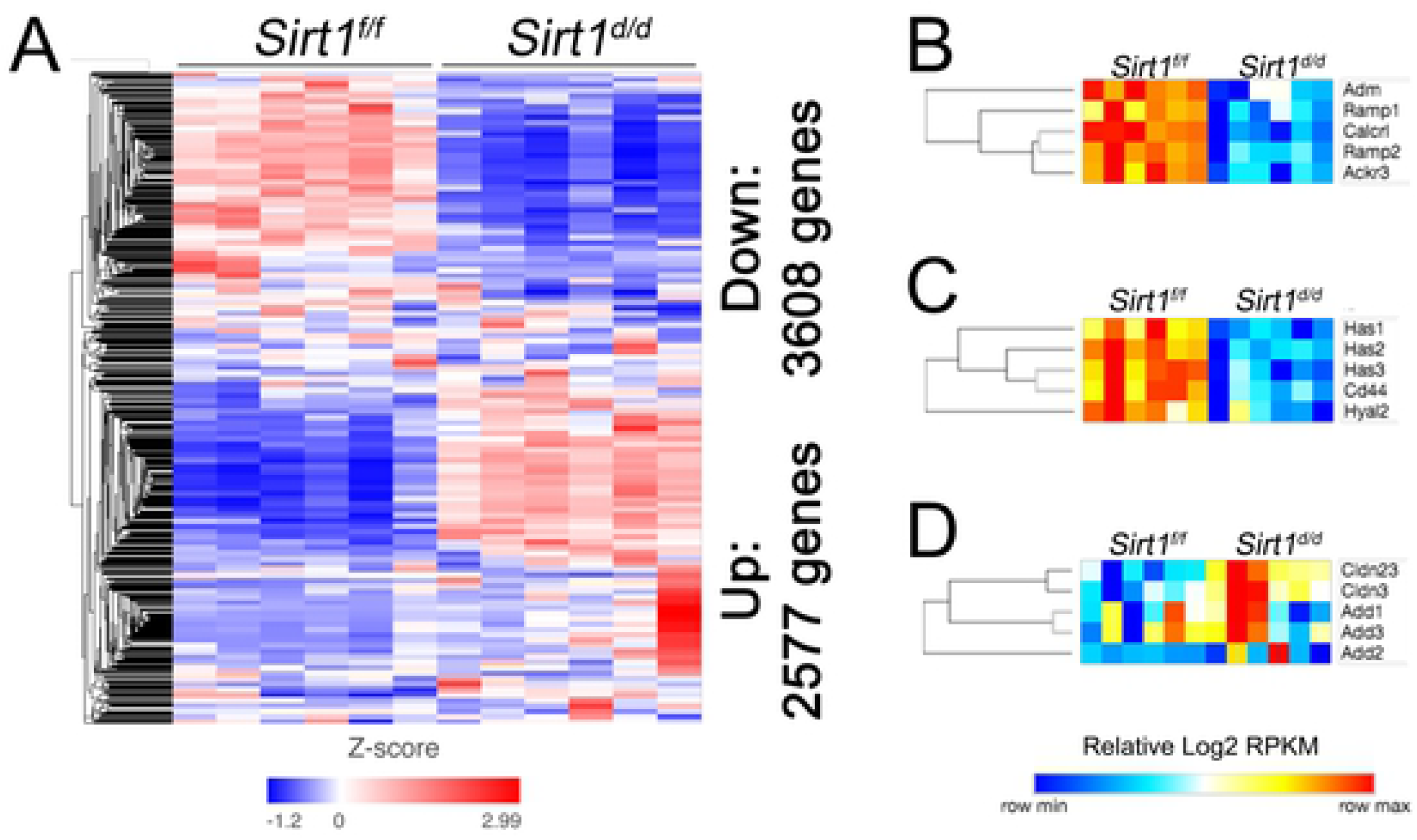
SIRT1-dependent expression of uterine genes. **(A)** Hierarchical clustering heatmap of gene expression levels in *Sirt1^f/f^* and *Sirt1^d/d^* mouse uteri at GD 3.5 (n=6) as determined by RNA-seq. Differential gene expression analyses identified 6,185 uterine genes significantly regulated in the *Sirt1^d/d^* mice relative to *Sirt1^f/f^*. **(B-D)** Hierarchical clustering heatmaps of RNA-seq data showing expression levels (log_2_ RPKM) of genes related to the adrenomedullin (ADM) signaling pathway (B), hyaluronic acid (HA) signaling (C), and formation of epithelial tight junctions (D) in *Sirt1^f/f^* and *Sirt1^d/d^* mouse uteri at GD 3.5 (n=6).

**S4 Fig.**
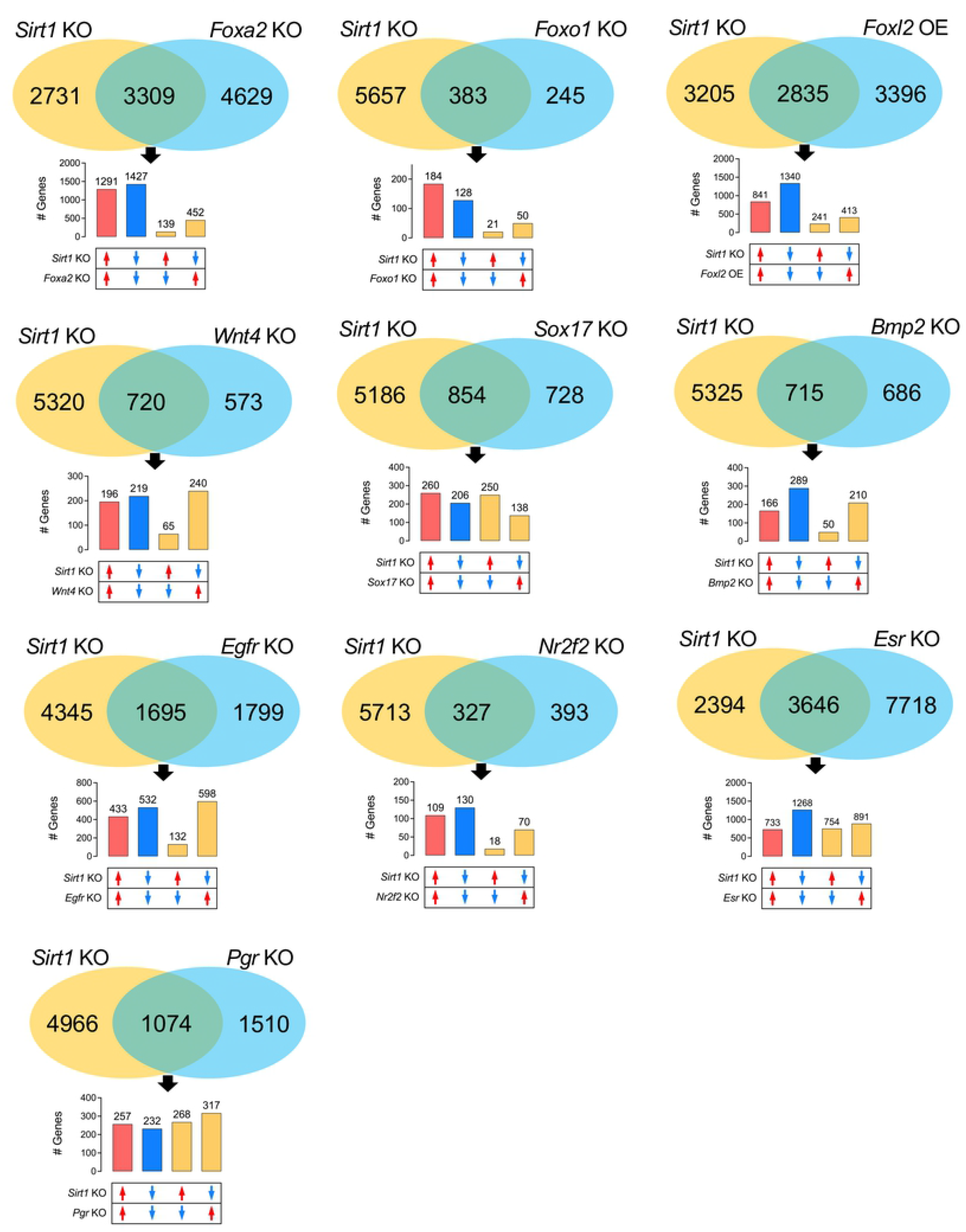
Overlaps and correlations of *Sirt1* KO (GD 3.5) with *Foxa2* KO (GD 3.5), *Foxo1* KO (PPD 4.5), *Foxl2* OE (diestrus), *Wnt4* KO (GD 3.5), *Sox17* KO (GD 3.5), *Bmp2* KO (GD 3.5), *Egfr* KO (GD 3.5), *Nr2f2* KO (GD 3.5), *Esr* KO (ovariectomized mice 24 h after E2 injection) and *Pgr* KO (GD 3.5) transcriptomes in mouse uteri. KO, knockout; OE, overexpression; GD, gestational day; PPD, pseudopregnant day; DD, decidual day.

## Supplemental#Tables

**S1 Table. List of differentially expressed genes in *Sirt1^d/d^* mouse uteri compared with *Sirt1^f/f^* mouse uteri at GD 3.5.**

**S2 Table. Functional annotation of altered transcriptome in *Sirt1*-deficient mouse uteri at GD 3.5.**

**S3 Table. Upstream regulators of altered transcriptome in *Sirt1*-deficient mouse uteri at GD 3.5.**

**S4 Table. List of commonly regulated genes between *Sirt1*-deficient and aged mouse uteri at GD 3.5.**

**S5 Table. Information for primers and probes. S6 Table. Antibody information.**

